# Single residue changes to the histone core catalyze neofunctionalization and impose fitness trade-offs

**DOI:** 10.1101/2024.05.10.593535

**Authors:** Zachary H. Harvey, Kathryn M. Stevens, Jian Yi Kok, Akihisa Osakabe, Jiaying Liu, Tobias Warnecke, Frédéric Berger

**Affiliations:** Gregor Mendel Institute, Austrian Academy of Sciences, Vienna, Austria; MRC Laboratory of Medical Sciences, London, United Kingdom; Institute of Clinical Sciences, Imperial College London, London, United Kingdom; Department of Biological Sciences, Graduate School of Science, The University of Tokyo, Tokyo, Japan; Trinity College, Oxford, United Kingdom; Department of Biochemistry, University of Oxford, Oxford, United Kingdom

## Abstract

Histones are among the most conserved proteins in the eukaryotic genome, and their function is thought to be largely invariant across species. Here, we tested this assumption, examining over a billion years of the essential histone H2A.Z’s evolution in a single synthetic host. We identify single residue substitutions within the H2A.Z core domain that led to its neofunctionalization. Such H2A.Z neomorphs are distinct by their ability to directly interact with the transcription apparatus, rewiring gene expression genome-wide by tuning transcription processivity. Our results reveal that even changes of single residues within the histones core domain can transform their function, catalysing the rapid emergence of phenotypic diversity by directly imposing both fitness opportunities and costs. We propose that the entire histone sequence has the potential to evolve new regulatory relationships, providing a framework to understand the mechanistic underpinnings of disease-associated histone mutations.

## Main Text

Histone proteins are nearly invariant across eukaryotes,^1^ constrained by their central position in genome biology from DNA packaging^2^ and repair^3^ to transcription.^4–6^ Indicative of this, histone mutation at the scale of even single residues is costly: so-called oncohistone mutations are potent drivers of human disease,^7–10^ and species barriers among histone proteins can arise with only a handful of changes.^11^ Yet, diverse histone forms have evolved,^1,12^ and are central to regulating fundamental biological processes such as development,^13,14^ suggesting changes in histones sequence also afford adaptive opportunities. In this frame, it remains unknown how histones can navigate their constraints to acquire new functions with only small sequence changes, and what forces balance such changes. We examined over a billion years of the essential histone H2A.Z’s evolution using the fission yeast *Schizosaccharomyces pombe* as a synthetic chassis.^11,15–17^ Single substitutions within the histone core domain loop 2 (L2) region were sufficient to drive emergent transcriptional properties, in turn reshaping the cell’s response to abiotic stress. Such neomorphic H2A.Zs often acquired a protein–protein interaction between the L2 and transcription elongation factor Spt6, driving its recruitment to chromatin and increasing polymerase processivity *in vivo*. Together, our findings demonstrate how histones can, through even single residue changes, acquire new functions. Further, they highlight the potential for neofunctionalization within the histone core domain. These molecular insights offer a window into the events driving chromatin’s evolution and identify potential new avenues of investigation to understand the molecular pathology of the thousands of oncohistone mutations^10^ yet to be characterized.

### Diverse H2A.Zs from Extant Eukaryotes Encode Emergent Functions

To investigate the mechanisms by which histones can acquire new functions, we focused on the essential histone H2A.Z. H2A.Z is among the most conserved histone proteins encoded by eukaryotic genomes,^1^ where it has been linked to transcription regulation.^18–20^ We first constructed a database of H2A.Z sequences from the EukProt database^21^ (Extended Data Figs. 1, 2a, & 3; Table S1; Data File S1; Methods). As previously suggested from analyses of smaller datasets,^1,22^ we found that H2A.Z forms a monophyletic clade that spans eukaryotic diversity (Extended Data Figs. 2b & 3a). Importantly, we found H2A.Z orthologs in deep-branching clades like Metamonada that were previously thought to lack H2A.Z (Extended Data Fig. 2b).^23^ Thus, the phylogenetic distribution points to H2A.Z being present in the last eukaryotic common ancestor (LECA), whilst being broadly retained throughout the evolution of eukaryotes.

We selected nine H2A.Z sequences from across the phylogenetic tree (Fig. 1a; Extended Data Figs. 2b & 4a, Data File S2), chosen to maximize the number of eukaryotic supergroups represented and their total sequence diversity. Although they span more than a billion years of evolution, representative H2A.Zs were ∼65–90% identical in their core domain (excluding the highly divergent N- and C-terminal tail regions; Extended Data Fig. 4b), highlighting their high degree of conservation.

**Figure 1.**
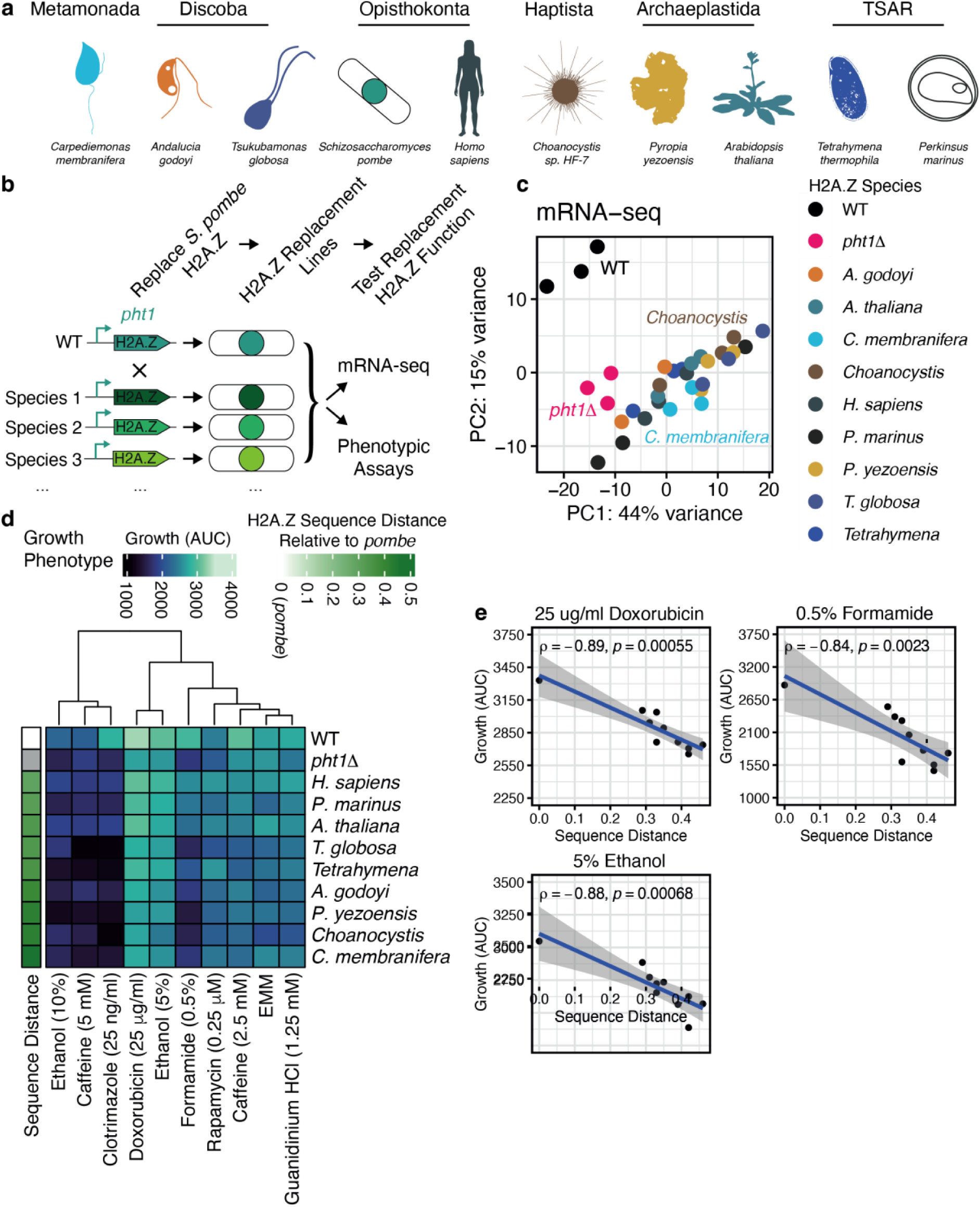
Diverse H2A.Zs from Extant Eukaryotes Encode Emergent Functions. **a,** Supergroup diversity of chosen lineages for synthetic replacement experiments detailed in (b). For each supergroup, representative species is written below. **b,** Experimental scheme describing the replacement of endogenous *S. pombe pht1* by representative H2A.Zs taken from the H2A phylogeny (Extended Data Fig. 2b, 3a). **c**, Principal component analysis (PCA) for the top 500 variable genes from mRNA-seq data generated for H2A.Z replacement lines. **d**, Growth phenotypes for H2A.Z replacement lines in various stress conditions, annotated with the sequence distance of each H2A.Z from that of *S. pombe* Pht1 (which has a sequence distance of zero). Plotted values are the mean of four biological replicates. Growth is quantified as the area under the curve (AUC) from respective growth curves. H2A.Zs are annotated with their sequence distance relative to *S. pombe* Pht1, with zero being Pht1 and larger numbers indicating greater divergence. **e**, Example Spearman’s correlations between the growth from **c** with sequence distance of each H2A.Z relative to WT *S. pombe*.

To test whether transcription is impacted by sequence diversity in these nine representative H2A.Z proteins, we generated yeast lines replacing the sole endogenous H2A.Z gene (*pht1*) with one of these orthologs (Fig. 1b). We then assessed global transcriptome changes using strand-specific mRNA-seq using WT and an H2A.Z deletion mutant (*pht1*Δ) as references. Whereas *pht1*Δ cells exhibited modest transcriptional changes, affecting ∼10% of detected transcripts (Extended Data Fig. 4c), H2A.Z replacement lines showed changes in ∼12– 23% of detected transcripts when compared with WT (Extended Data Fig. 4c). Further, principal component analysis of their transcriptomes revealed that transcriptomes of H2A.Z replacement lines were distinct from both WT and *pht1*Δ (Fig. 1c). These data indicate that H2A.Zs from different species are not graded loss-of-function alleles, rather giving rise to sequence-specific emergent transcriptional states.

We next asked how transcriptional variation in H2A.Z from different species impacts phenotype. To do this, we assessed the growth of H2A.Z replacement lines under stress conditions targeting metabolism (rapamycin^24^), protein folding (low-dose guanidinium hydrochloride^25^), cell wall biogenesis (the anti-fungal drug clotrimazole^26^), fermentation (ethanol^27^), or chromatin and genome function (doxorubicin,^28^ formamide^29^, and caffeine^30^) (Fig. 1d). Surprisingly, H2A.Z replacement lines generally grew more slowly than *pht1*Δ cells (Fig. 1d; Extended Data Fig. 5a–c), with sequence divergence from *S. pombe* Pht1 strongly inversely correlating with growth (Fig. 1e, Extended Data Fig. 5d). By contrast, we saw no strong correlation between the level of H2A.Z expression and growth (Extended Data Fig. 5d–e), indicating that these effects were not attributable to differential expression of H2A.Z. Together, these data link primary sequence variation in H2A.Z from different species to emergent functional variation, suggesting they are neomorphs.

### Single Residue H2A.Z Core Domain L2 Substitutions Are Sufficient for Neofunctionalization

We next asked which H2A.Z sequence changes could drive neofunctionalization. Recent reports have suggested that the histone core domain, particularly the solvent-accessible^31^ M6 region surrounding the loop 2 (L2) is essential for H2A.Z function.^32,33^ Consistent with this, we found that the L2 alone was sufficient to complement *pht1*Δ (Extended Data Fig. 6a–e), and it was among the most conserved regions in H2A.Z across all sequences in our dataset (Fig. 2a; Extended Data Fig. 3). Further, L2 sequence divergence relative to WT *S. pombe* Pht1 was the best correlated with the transcriptional changes observed in full-length H2A.Z replacement experiments (ρ = 0.63, *p* < 10^-3^, Fig. 2b; Extended Data Fig. 6f), suggesting it alone is a strong predictor of neofunctionalization of H2A.Z impact on transcriptomes.

**Figure 2.**
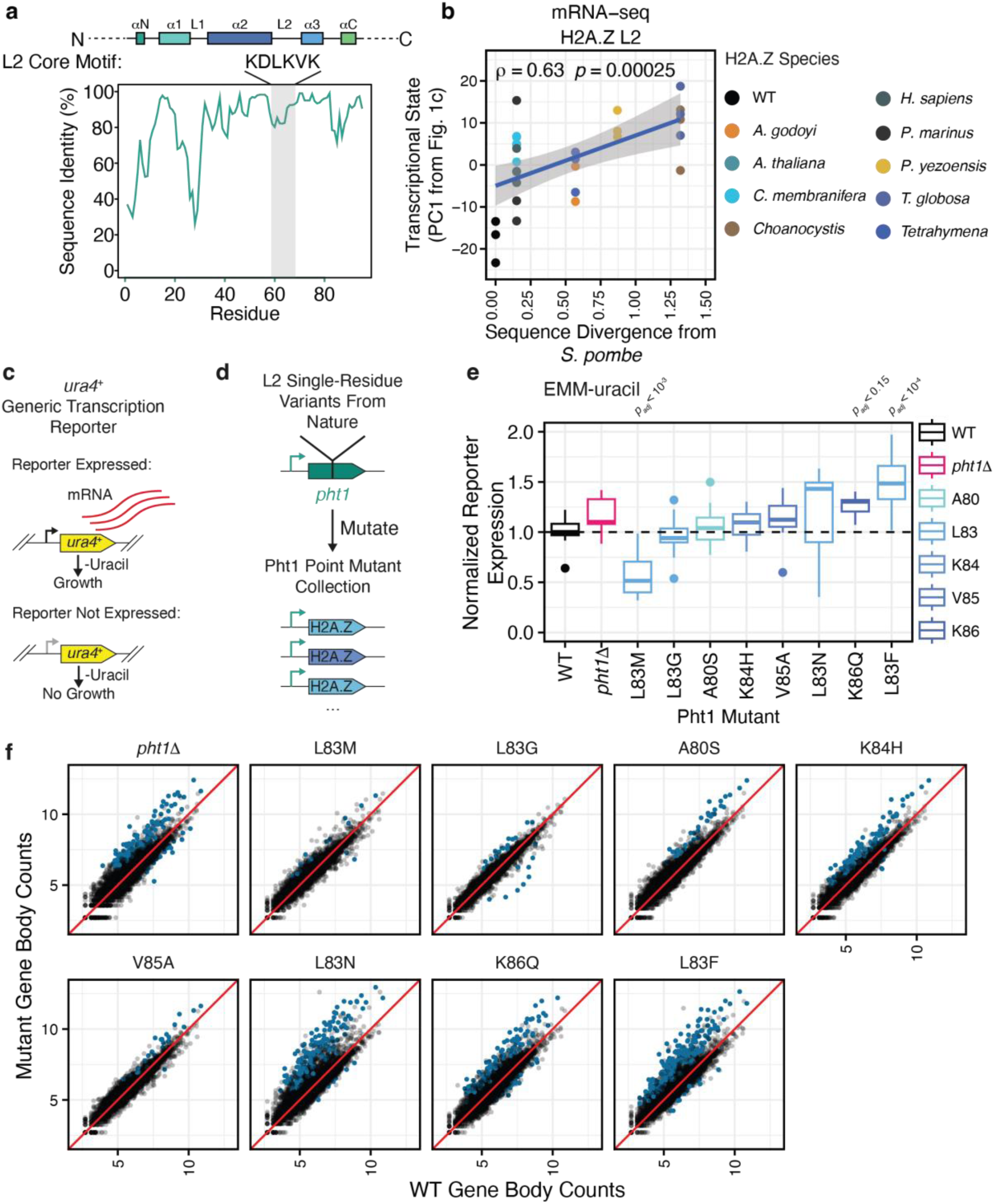
Single Residue H2A.Z Core Domain L2 Substitutions Are Sufficient for Neofunctionalization. **a**, Mean pairwise sequence identity of H2A.Z and H2A.X residues across eukaryotes, averaged using a 3-amino acid sliding window. The L2 region is highlighted in grey on the histone secondary structure, with the core domain we identify indicated. Dashed lines indicate the N- and C-terminal tails that were not included in this analysis due to variable length and resulting gappy alignment. **b**, Spearman’s correlation analysis between the sequence divergence from *S. pombe* Pht1 L2 and principal component 1 from mRNA-seq analysis from Fig. 1b. In this analysis, the *S. pombe* Pht1 sequence corresponds to a value of zero, with a larger number corresponding to a greater divergence. See also Extended Data Fig. 5e. **c,** Experimental scheme explaining the *ura4*^+^ reporter gene. **d,** Experimental scheme describing the generation of H2A.Z point mutants. **e,** WT-normalized growth of H2A.Z point mutant strains bearing the transcriptional reporter from **c** in EMM-Uracil. The median of WT is represented with a dashed line. Each Tukey boxplot represents 12 biological replicates, adjusted *p*-values (*p_adj_*) are calculated using a two-way ANOVA followed by a post-hoc Tukey test. **e,** Differential count analysis for gene body qPRO-seq signal calculated using DESeq2. Red line, diagonal (Y=X) line drawn to aid in interpretation. Statistically significant (*p_adj_*<0.1) changes are indicated in blue.

To test whether H2A.Z L2 sequence variation was sufficient to propel neofunctionalization, we first constructed a transcription reporter for H2A.Z using the uracil biosynthesis gene *ura4* (Extended Data Fig. 7a–d). In this assay, the amount of transcription of *ura4* is directly related to the amount of growth in media lacking uracil (Fig. 2c), allowing rapid genetic interrogation of the importance of each individual residue in the L2. We thus tested whether individual substitutions within the L2s from representative species (Extended Data Fig. 4a) were sufficient to change expression of the reporter relative to either WT or *pht1*Δ cells (Fig. 2d). Whereas *pht1*Δ cells had modestly increased reporter expression (Fig. 2e), L2 point variants were generally stronger. Indeed, half of L2 substitutions exhibited a strong increase (by ∼1.3–1.5 fold) or decrease (by ∼2 fold) in reporter expression (Fig. 2e). Importantly, Pht1 L2 variants did not have general fitness defects (Extended Data Fig. 7e). Further, Pht1 L2 variant incorporation into chromatin was required for them to impact to transcription because, in absence of the H2A.Z-depositing Swr1 complex (*swc2*Δ background^34,35^), all differences in L2 variants relative to WT, except for L83M, were abrogated (Extended Data Fig. 7f). Finally, we checked whether Pht1 L2 variants were sensitive or resistant to the transcription inhibitor MPA.^36^ In this assay, Pht1 L2 variants were either sensitive (L83M) or robust (*pht1*Δ, L83N, K86Q, and L83F) to treatment (Extended Data Fig. 7g), supporting their impact on transcription in reporter assays. Together, these data establish that even single-residue H2A.Z L2 variants can generate transcriptional states distinct from either WT or *pht1*Δ and are therefore transcriptional neomorphs.

Having established a genetic link between L2 neomorphs and transcription, we next examined their genomic impact. Consistent with results of our reporter assays, differential expression analysis of gene body-located nascent transcripts in WT and *pht1*Δ lines, a measure of active transcription obtained by qPRO-seq,^37,38^ showed a modest increase in transcription in *pht1*Δ cells (Fig. 2f). Further, the strongest mutants in our reporter assays exhibited ∼50% greater increase in transcription than in *pht1*Δ (Fig. 2f). Interestingly, the single neomorph that did not concur with our reporter assays, L83M, nevertheless had decreased *ura4* mRNA levels (Extended Data Fig. 7c), and was the only tested variant to impact nucleosome stability (Extended Data Fig. 7h), suggesting it acts indirectly on transcription. To measure the impact of enrichment of the diverse forms of Pht1 we used ChIP-seq to obtain their genomic profiles. Whereas overall levels of Pht1 were similar across L2 neomorphs, we did note a loss of TSS enrichment in all neomorphs (Extended Data Fig. 8a). However, such changes were not correlated with transcriptional output genome-wide (Extended Data Fig. 8b). Taken with L2 neomorphs being genetically downstream of their deposition by Swr1-C (Extended Data Fig. 7f), these observations suggest that changes to Pht1 genomic distribution are downstream of their impact on transcription. Together, these data establish that sequence variation in the core L2 region of H2A.Z is sufficient to transform its transcriptional impact on the genome.

### H2A.Z Neomorphs Rewire Its Interaction with The Transcription Apparatus

We next investigated whether the impact of H2A.Z neomorphs on the transcription apparatus was direct. In the context of the nucleosome, the L2 region of H2A.Z is solvent accessible,^39^ suggesting it could form protein–protein interactions. From our qPRO-seq analysis, we observed the strongest effects specifically on transcription elongation, as indicated both by the large increase in gene body nascent transcripts relative to WT (Fig. 2f) and resistance to the inhibitor MPA, which preferentially inhibits transcription elongation^36^ (Extended Data Fig. 7g). Taking these together, we wondered whether L2 neomorphs might gain an interactor involved in transcription elongation. Previous reports have established an antagonistic relationship between H2A.Z^40^ and the transcription elongation factor Spt6, which both regulates transcription processivity.^41,42^ We wondered whether L2 neomorphs might modify this relationship. To ascertain whether this is a possibility, we first performed an unbiased AlphaFold2 Multimer^43^ interaction screen for WT and single-residue variants of the L2 core motif from our database of H2A.Z sequences (Extended Data Fig. 3, Data File S1) against the *S. pombe* nuclear proteome.^44,45^ We then examined which predicted interactions varied with L2 sequence (Extended Data Fig. 9a–b). Spt6 emerged as amongst the highest variance interactors in this dataset (Extended Data Fig. 9c; Supplementary Table S3, S4), often gaining a strong interaction score relative to WT Pht1. Further, it was the only potential interactor that combined highly robust folding models across multiple simulations (Extended Data Fig. 9d) and predicted structures closely matching experimentally derived structures (Extended Data Fig. 9e).

To test these *in silico* predictions, we first established an *in vitro* pulldown assay using recombinantly expressed and purified Spt6, finding an interaction between H2A.Z and Spt6 was possible either with full-length Pht1 (Extended Data Fig. 9f–h), or a peptide corresponding only to the L2 (Fig. 3a–b). To test whether L2 sequence variation could impact the Spt6–L2 interaction, we next synthesized selected H2A.Z L2s from different species in our test set (Extended Data Fig. 4a), as well as a pair of the two most extreme single-residue neomorphs from our transcription assays (Fig. 2d–e) and assayed their binding to Spt6 (Fig. 3a–b). Variation in Pht1 L2 sequence was sufficient to influence the amount bound to Spt6 (Fig. 3b). These data establish that the L2 region alone is sufficient to account for the interaction of Pht1 with Spt6.

**Figure 3.**
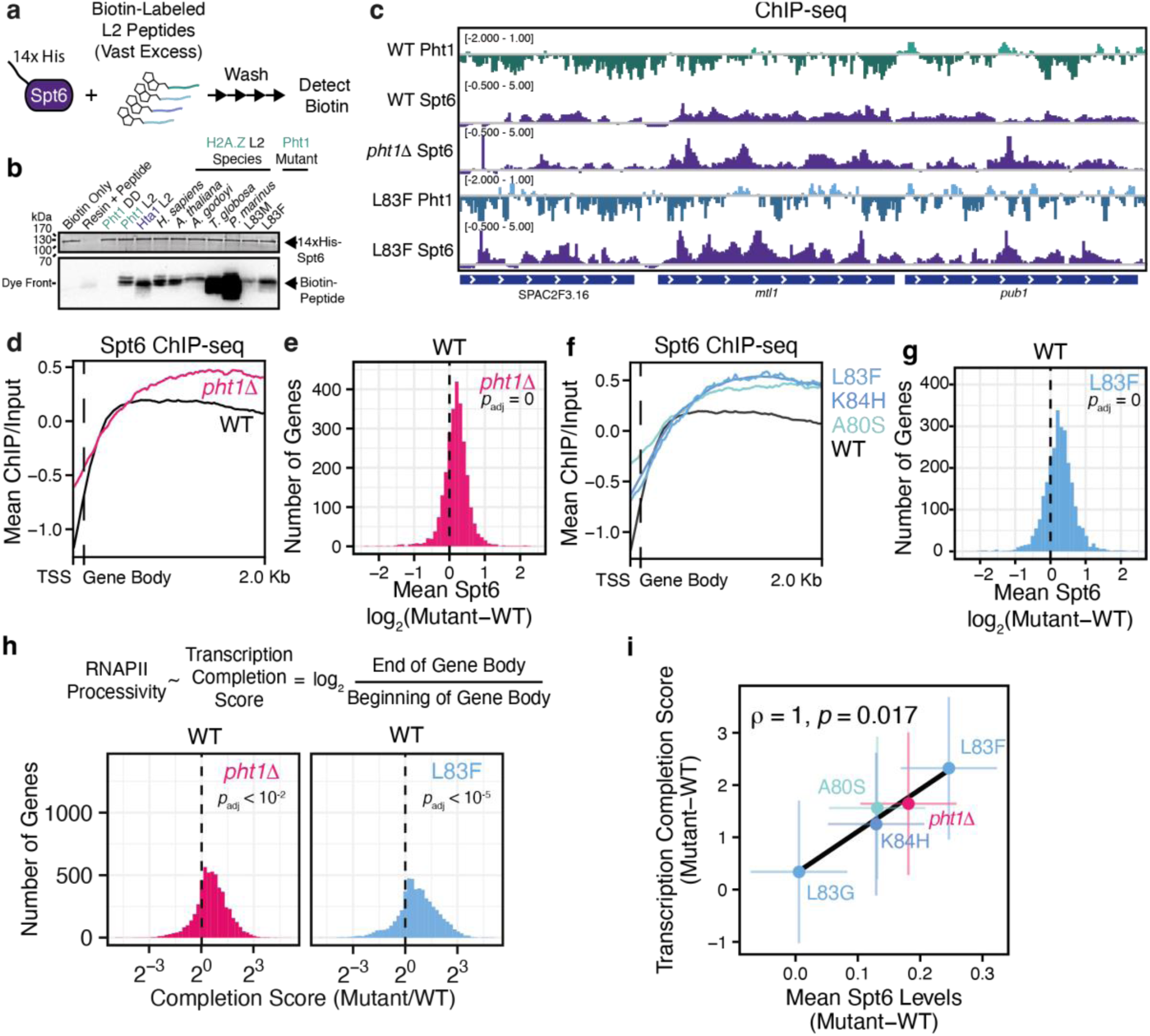
H2A.Z Neomorphs Rewire Its Interaction with The Transcription Apparatus. **a,** Experimental scheme showing the pulldown in **b** of biotinylated peptides corresponding to either WT or L2 neomorphs against WT *S. pombe* 14xHis-Spt6(282-1159) immobilized on Ni-NTA resin. Detection was done by streptavidin-HRP conjugate. Representative of five experiments, each with similar results. Hta1 L2 and Pht1 docking domain (DD) are included as positive and negative controls, respectively. **c,** Example ChIP-seq tracks for Pht1, Pht1 neomorph, and WT-Spt6 in the indicated Pht1 mutant background. Plotted is the log_2_ transformed, input-normalized signal. **d–g,** Analysis of genome-wide Spt6 levels within gene bodies for indicated *pht1* mutants. Plotted is the mean input-normalized Spt6 signal for at least two biological replicates either as a mean profile for **d & f** or histogram with the reference value for WT indicated by a dashed line in **e & g**. Adjusted *p*-values are from a post-hoc Tukey test of a 2-way ANOVA. **h,** Transcription completion scores calculated using the indicated formula from qPRO-seq data in the respective mutants, expressed as the change relative to WT. Adjusted *p*-value from a post-hoc Tukey test of a 2-way ANOVA. **i,** Correlation analysis of the average change in transcription completion score relative to WT with respect to the change in Spt6 levels, also with respect to WT, for the indicated Pht1 neomorphs. Correlation coefficient is Spearman’s rho, and average values for each mutant are bounded by the 95% confidence interval.

We next investigated the consequences of the interaction between L2 neomorphs and Spt6 on transcription elongation *in vivo*. To do this, we performed ChIP-seq against Spt6 in either WT or L2 neomorphs (Fig. 3c), comparing the results to transcriptional output as measured by qPRO-seq. Generally, *pht1* mutants lead to increased Spt6 levels genome wide (Fig. 3c–g; Extended Data Fig. 10a, except L83G), with Spt6 levels increasing moderately in *pht1*Δ cells relative to WT (Fig. 3d–e), and extremely in L2 neomorphs (Fig. 3f–g; Extended Data Fig. 10a). Given Spt6’s link to stimulating RNAPII processivity,^41,42^ we next asked whether increased Spt6 enrichment in chromatin was associated with increased processivity. To do this, we used qPRO-seq data to calculate the ratio of nascent transcripts at the end of a gene relative to those near the transcription start site (TSS), a metric known as the completion score that reflects changes in RNAPII processivity.^41^ Increased Spt6 levels associated with *pht1* mutants generally positively correlated with higher transcription completion scores (Fig. 3h; Extended Data Fig. 10b), and there was an essentially perfect correlation between the global increase in Spt6 levels for a given L2 neomorph and global processivity increases (ρ = 1, *p* < 0.02; Fig. 3i). Together, these results establish a direct link between L2 neomorphs acquisition of a protein–protein interaction with Spt6, its recruitment to chromatin, and transcription stimulation.

### L2 Neomorphs Balance Phenotypic Opportunities and Costs Through Transcriptional Regulation

Finally, we considered why, despite apparent regulatory potential, L2 neomorphs are only rarely found in naturally occurring sequences. To do this, we expanded our analysis beyond natural variation by performing deep scanning mutagenesis^48^ of the 7-residue H2A.Z L2 core motif. Briefly, we generated a library of all 133 possible single residue substitutions using a scarless CRISPR-based strategy^49^ (Fig. 4a). Remarkably, this yielded a wide range of transcriptional states (Fig. 4b; Extended Data Fig. 11a), with each position within the L2 having a unique profile (Fig. 4c). Indeed, although many positions could lead to similar transcriptional output, it was rare that the same amino acid substitution led to the same outcome at different positions. For example, whereas arginine was activating at position L83, it was repressive at position K86. The only exception was proline, which had a position-independent stimulating effect (Extended Data Fig. 12a). Together these results establish that both position and amino acid identity in the L2 are integrated to define a broad spectrum of transcriptional outputs.

**Figure 4.**
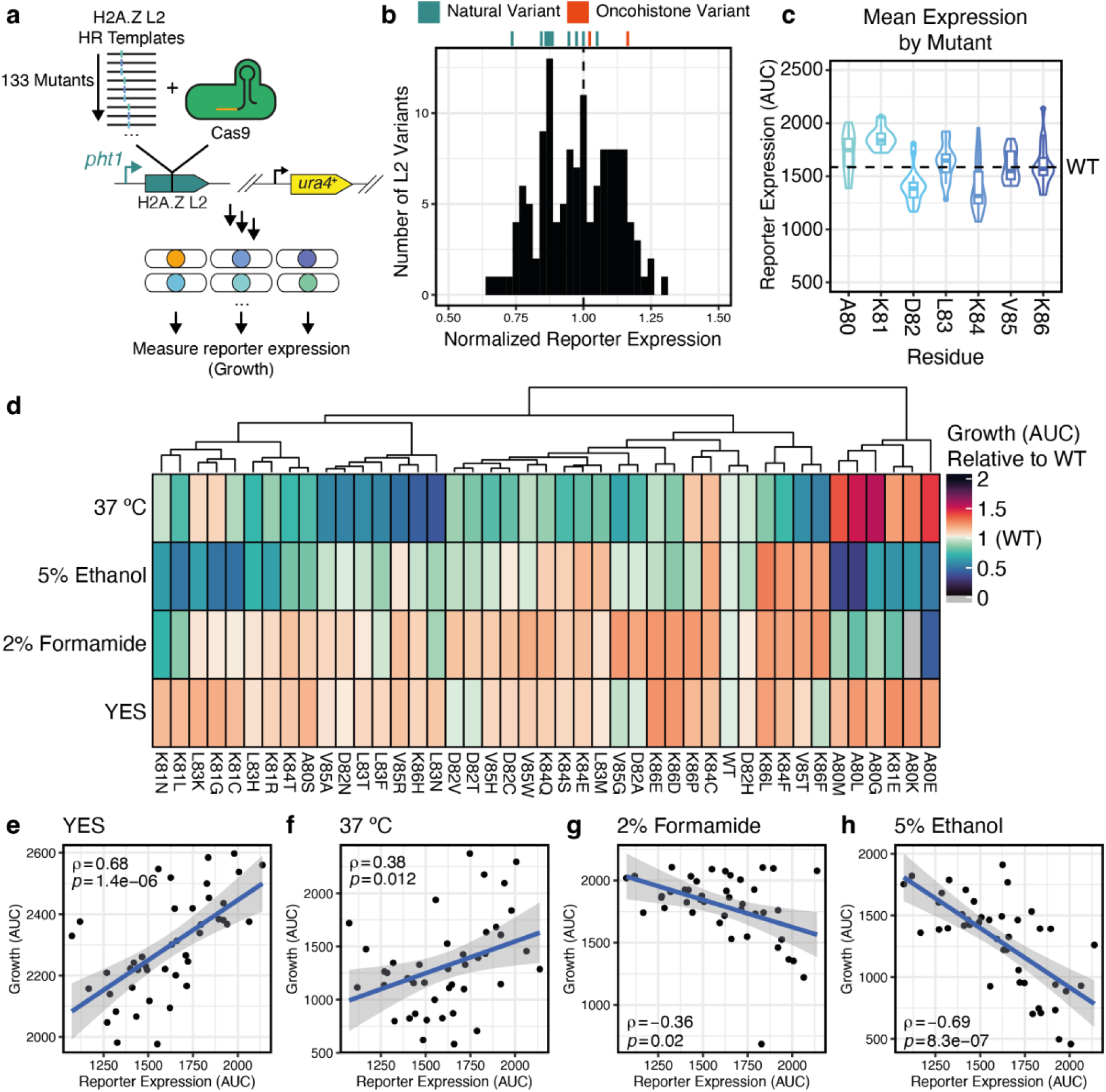
L2 Neomorphs Balance Phenotypic Opportunities and Costs Through Transcriptional Regulation. **a,** Schematic of CRISPR strategy for deep scanning mutagenesis of the endogenous Pht1 L2. **b,** Histogram for the mean *ura4* reporter expression for all L2 neomorphs. Histogram is based on the mean of 6 biological replicates, and bins including either L2 sequence variants (at Hamming Distance = 1 relative to WT S. pombe Pht1 L2) or known oncohistone variants^10^ labeled above. **c,** Violin plots for the data from **b** broken down by individual positions within the L2. **d,** Growth relative to WT for Pht1 L2 neomorphs in *S. pombe* across four conditions. Growth is calculated as the area under the growth curve for 6 biological replicates. **e–h,** Spearman’s correlation analysis for mutants from **d** with their transcriptional reporter output from **b**.

Although many activities are in principle possible, we wondered whether nature retains certain variants over others. We identified only 12 L2 neomorphs that occur in nature (Hamming distance=1 relative to WT *S. pombe* Pht1; Data File S1), as well as a pair of oncohistone mutants (Fig. 4b).^10^ Whereas oncohistone mutants were neutral-to-activating side of the distribution, natural L2 sequences were notably depleted from this side of the distribution (*p* < 0.01; Extended Data Fig 11b–c). This led us to ask why activating variants, which we found were associated with gaining a direct protein–protein interaction with transcription, were not favoured in natural sequences. Although phenotypically neutral under defined conditions (Extended Data Fig. 7e), we found that transcriptionally-activating neomorphs had a growth advantage under nutrient-replete media (YES) conditions (ρ = 0.69, *p* < 10^-5^; Fig. 4d–e). Notably, such metabolic conditions parallel those experienced by early-stage tumour growth.^50^ By contrast, either a mild heat,^51^ RNA metabolism,^29^ or ethanol stress eliminated or even reversed this relationship (ρ = −0.68, *p* < 10^-6^; Fig. 4d,f–h). Thus, whereas under relatively favourable conditions, L2 neomorphs acquiring a direct connection to transcription offers a strong phenotypic advantage, under stressful conditions, this relationship is neutral or even detrimental. Together, these data suggest that L2 neomorphs can drive neofunctionalization of histones and influence organismal fitness in different environmental conditions.

## Discussion

Here we describe how the essential histone H2A.Z neofunctionalizes, providing evidence that one of the fundamental mechanisms by which histones can acquire new functions is by establishing protein–protein interactions between the histone core domain and the transcriptional apparatus. Examining naturally occurring sequences in a synthetic system, we find that even single amino acid substitutions within the 7-amino-acid long histone core domain loop 2 (L2) of H2A.Z is a powerful source of new regulatory potential. Indeed, H2A.Z L2 neomorphs often gain an interaction with transcription elongation factor Spt6, driving its recruitment to chromatin and increase transcription processivity. Finally, we detail the consequences of the L2 neomorph–Spt6 regulatory relationship, demonstrating that it presents both phenotypic opportunities, affording phenotypic advantage under favourable conditions, but also imposing a cost in stressful environments. Together, our results provide a mechanism whereby a single residue change in a conserved protein can rewire global gene expression and precipitate phenotypic diversification.

The core histone fold domain has long been known to be important for the interaction with DNA and the stability and structure of the nucleosome.^20,52–59^ Our findings reveal an additional layer of potential regulation by highlighting an expanded landscape of protein– protein interactions between the histone core and the transcription apparatus. We show that even the smallest possible unit of change to amino acid sequence can lead to a complete change in a histone’s function. This highlights the evolutionary power of ultra-conserved proteins for functional innovation, suggesting that functional and phenotypic change do not always require large-scale changes to achieve emergent activities. In this frame, we anticipate that histones, and by extension nucleosomes, encode a diverse repertoire of regulatory mechanisms that yet await description.

Finally, our results posit a new framework to understand oncohistone mutations associated with various cancers.^10^ Although there have already been efforts to understand the mechanistic basis of oncohistones,^60–62^ thousands remain uncharacterized.^10^ We find that many independent mutants converge on similar mechanisms, it highlights the possibility that, whereas most oncohistone mutants are associated with specific types of cancers, patients, and even tumors, they may converge on similar mechanisms. Our results outline an experimental approach whereby gain-of-function, emergent effects of such mutants can be identified and dissected, opening the door to new potential therapeutic avenues.

## Methods

### Histone Dataset Construction

To investigate H2A.Z sequence diversity across eukaryotes, we sought to identify H2A.Z orthologs using a phylogenetic approach. First, to identify proteins homologous to a specific core histone (i.e. to H2A rather than H2B, H3, or H4), we used HHMer^63^ to build specific HMM models (command = hmmbuild --amino) for each of the four core histones from curated histone sequences in HistoneDB 2.0.^64^ The resulting models retrieve all except one histone variant in HistoneDB (H2A.P from *Heterocephalus glaber,* annotated as ‘Huntingtin-interacting protein M’) and hit the wrong core histone in only three instances. This suggests that the models are both sensitive and specific and can be used to discriminate different core histones from each other with reasonably accuracy. We therefore proceeded to search for hits against each model (hmmsearch –noali) in the EukProt protein database (v2, accessed on 01-12-2020),^21^ retaining only hits with an E-value < 0.001. With this filter, the probability that a given histone is hit by more than one core histone-specific HMM is relatively low (Extended Data Fig. 1a).

To determine whether separate HMM models for each H2A *variant* could be used to identify H2A *variants* by homologous search alone, we build models in the same manner as above (using curated H2A variants from HistoneDB) and then searched EukProt and HistoneDB 2.0. Variant HMM models hit the correct HistoneDB H2A variant in most instances, but a given variant is often also hit by additional variant models (Extended Data Fig. 1b). Even though E-values can be used to identify the correct variant in most instances (Extended Data Fig. 1c), we consider this level of specificity insufficient for robust identification of H2A variants and therefore proceeded with homologs identified by the initial H2A HMM as the starting point for tree building. Histones also hit by another core histone HMM were only retained if the H2A model returned the best E-value.

### Tree Building

To facilitate tree building, we only retained putative H2A orthologs shorter than 400 amino acids and only considered H2As from species with hits against all four core histone HMMs (568 out of 742 species). Some species, including metazoa and plants, encode many more H2A paralogs than others. To balance the dataset, we retained a maximum of five species per supergroup, eliminating species semi-randomly (prioritising coverage of different Eukprot lineages). The resulting dataset contains 839 putative H2As from 127 species (Extended Data Fig. 3).

These putative H2A sequences were aligned using MAFFT (command= mafft -- localpair --maxiterate 1000)^65^. ModelTest-NG^66^ was used to determine the best fit model of evolution to be used for tree building (WAG+G4), which was done using RAxML-NG (command = raxml-ng --msa --model WAG+G4 --seed 2)^67^. Sequences associated with long branches (Extended Data Fig. 3a) were manually inspected but not removed as their presence does not affect conclusions. Trees were plotted using iTOL.^68^

### Yeast Culture, Strain Manipulation, and Phenotypic Assays

All strains used in this study are listed in Supplementary Table S5. Yeast (*S. pombe* 972 *h*-) were cultured under standard conditions (32 °C). Unless otherwise stated, media used were either EMM2 (Edinburgh Minimal Media)^69^ or YES (Yeast Extract with Supplements).^70^

*S. pombe strains* were manipulated either using one-step PCR-based HR mutagenesis^71^ or the SpEDIT CRISPR editing platform.^49^ For both, heat-shock was used to transform chemically-competent parent strains.^72^ Donor templates for homologous recombination were prepared using either Gibson assembly^73^ using reagents supplied by the IMP Molecular Biology Service combined with site directed mutagenesis, or by direct synthesis from commercial vendors (IDT, Genscript). For commercially synthesized HR templates, sequences were codon optimized according to *S. pombe* codon usage using Benchling (https://benchling.com). In all experiments, WT control transformations were performed such that synonymous mutations were introduced to delete CRISPR gRNA target sites. All manipulations were verified by PCR genotyping and where appropriate sequencing of the relevant locus.

To construct the *ura4*-based reporter, a construct (PZH_352) consisting of homology to the 5**′** of the *ura4*-D18 locus,^74^ the selectable marker kanMX6,^71^ a 1 kb fragment corresponding to the weak promoter of *Arabidopsis thaliana ACT2* amplified from gDNA extracted from col-0, and the *S. pombe ura4*^+^ gene were assembled by Gibson. The *ura4-D18* locus was then replaced with the reporter via one-step PCR followed by homologous recombination.^71^ Correct integration was verified by PCR genotyping, as was accumulation of Pht1 over the reporter region by ChIP-seq (Extended Data Fig. 7b).

Phenotypic assays in this study were performed using a plate-based bulk growth assay. OD_600_ was monitored in 384-well plates (Nunc) continuously under the stress conditions indicated in figure legends at the standard growth temperature with shaking. Saturated cultures were sub-cultured 100-fold, and all assays were performed in either EMM complete, EMM without uracil (EMM-Ura), or YES. Data collection was performed on a BioTek Epoch2. Area under the curve (AUC) was calculated from these growth curves in R using the Growthcurver package (https://cran.r-project.org/web/packages/growthcurver/vignettes/Growthcurver-vignette.html). All subsequent data analysis was done in R.

### Antibody Generation, Protein Purification, and Analysis

A polyclonal antibody was raised in rabbit against *S. pombe* H2A.Z (Pht1) using a C-terminal-tail targeting peptide chosen based on predicted immunogenicity, and that it avoided the sequence domains of interest in the present study. Briefly, polyclonal serum was commercially produced (Eurogentec) using the peptide N-CLIRTKEKYPEEEEII-C. Antibodies were affinity purified using the same peptide. Briefly, SulfoLink agarose gel (Pierce Cat. No. 20401) was equilibrated in coupling buffer (50 mM Tris pH 8.5, 5 mM EDTA) before incubating with peptide for 90 min at RT. Afterwards, β-Mercapthoethanol was added and beads were incubated for 15 min at RT. Coupled beads were then washed three times in coupling buffer before adding serum diluted with 2 volumes of PBS with 0.2% tween-20. Beads and serum were then incubated together for three hours at RT with rotation. Beads were then collected in a disposable column and washed 6 times with PBS with 0.2% tween-20. To elute antibody, 750 µl of 0.1 M Glycine was added, incubated for 1 min, then allowed to flow through. Eluates were immediately quenched with 250 µl of 1 M K_2_HPO_4_. Elution was repeated nine additional times for a total of 10 fractions. Antibody-containing fractions were detected using Nanodrop spectrophotometer, pooling the highest concentration eluates. Antibodies were verified using *S. pombe* H2A.Z knockout strains (*pht1*Δ) for both Western blot and ChIP applications (Extended Data Fig. 7a), and aliquoted antibodies were stored at −70 °C.

Generally, recombinant proteins were produced by the Vienna BioCenter Core Facilities Protein Technologies Facility using constructs commercially synthesized (Genscript) in *Escherichia coli* BL21(DE3). *S. pombe* Hta1 and Pht1 were produced as full-length N-terminal GST fusions (PZH_668 and PZH_669). For *S. pombe* Spt6 a truncated version corresponding to residues 282–1159 (PZH_670) was produced as a 14xHis tagged fusion. Chimeric Spt6-*Hs*DLD (PZH_785) was produced by replacing the DLD of *S. pombe* Spt6 (residues 933–1061) with *E. coli* codon-optimized *Homo sapiens* Spt6 DLD (residues 1010– 1130) via Gibson assembly.

Expression of recombinant proteins was performed by autoinduction in TB media with 1.5% lactose and 50 µg/ml kanamycin for 24 hrs at 25 °C with shaking. Cells were lysed by sonication (Sonics VCX-750, 24-element or single-element microprobe: 3 mm, amplitude: 50%, time: 1 min, pulse-on: 1 s, pulse-off: 2 s) in the following buffer: 50 mM Tris-HCl pH 8.0, 500 mM NaCl, 5 mM DTT, Benzonase (2 μl per ml of Lysis buffer), 2 mM MgCl2, 0.2% (v/v) NP-40 and cOmplete protease inhibitor tablets (Roche, 1 tablet per 25 ml of lysis buffer).

*S. pombe* Pht1 and Hta1 were purified by glutathione affinity chromatography. Each supernatant was mixed with 100 μl of Glutathione Sepharose 4 FF (Cytiva Cat. No. 17513203, previously equilibrated with binding buffer A (50 mM Tris-HCl pH 8.0, 500 mM NaCl and 5 mM DTT) for 2h at 4 °C in a tube rotator. The beads were washed with 3 ml of binding buffer A. The protein was eluted with 400 μl of elution buffer A (50 mM Tris-HCl pH 8.0, 500 mM NaCl, 5 mM DTT and 10 mM reduced Glutathione). For GST-Hta1, the GST-elution was dialyzed against SP S buffer (25 mM HEPES pH 7.4, 100 mM NaCl, 1 mM DTT, 5% glycerol) for 18h in the cold room with constant mixing to adjust the conductivity for loading it onto 1mL HiTrap SP FF (Cytiva Cat. No. 17515701) column, pre-equilibrated with SP S buffer, at a flow rate of 1 ml/min. The column was then washed with SP S buffer until UV returned to baseline and the protein was eluted in a gradient to 50% of SP E buffer (25 mM HEPES pH 7.4, 1 M NaCl, 1 mM DTT, 5% glycerol) over 50 mL. Eluates were pooled and concentrated to a final volume of 1.5 mL using Vivaspin 20 MWCO 30 kDa (Sartorius Cat. No. VS2021). Concentrated pool eluates were then loaded onto a Superdex 200 16/60 (Cytiva Cat. No. 28989335) at a flow rate of 1 ml per minute previously equilibrated with SEC buffer (50 mM Tris pH 8.0, 150 mM NaCl, 1 mM DTT, 5% glycerol), where full-length protein eluted in the void volume.

For *S. pombe* Spt6 and Spt6-*Hs*DLD, purification was performed using immobilized metal affinity chromatography. Each supernatant was mixed with 500 μl of *Ni Sepharose 6 FF* (Cytiva Cat. No. 17531806, previously equilibrated with binding buffer B (50 mM Tris-HCl pH 8.0, 500 mM NaCl, 1 mM DTT and 20 mM imidazole)) for 2 hr at 4 °C in a tube rotator.

The beads were washed with 1.5 ml of binding buffer B and with 1.5 ml of wash buffer B (50 mM Tris-HCl pH 8.0, 500 mM NaCl, 1 mM DTT, 80 mM imidazole). The protein was eluted with 400 μl of elution buffer B (50 mM Tris-HCl pH 8.0, 500 mM NaCl, 1 mM DTT, 500 mM imidazole). All samples were dialysed against 1 l of dialysis buffer (25 mM Tris-HCl pH 8.0, 150 mM NaCl and 1 mM DTT) at 4 °C overnight. The dialysed samples were centrifuged at 4 °C and 21000xg for 5 min.

SDS-PAGE separation was performed using self-cast gels using standard methods. Gels were stained using colloidal blue reagents (MBS Blue) provided by the Molecular Biology Service. For Western blots, proteins were transferred using a standard wet protocol to a 0.2 µm nitrocellulose membrane (Cytiva Cat. No. 10600004), blocking in 5% non-fat dry milk (Maresi) dissolved in TBS^T^. Anti-Pht1 antibody was used at 1:1000 diluted in blocking buffer and detected using chemiluminescence following incubation with anti-rabbit HRP conjugates (Biorad Cat. No. 1706515), which were diluted 1:10,000 in blocking buffer. Imaging of gels and Western blots was performed using a Thermo Fisher iBright 1500 imaging system.

### Thermal Stability Assay of In Vitro Assembled Nucleosomes

Recombinant histones (both WT and mutants) were expressed in *E. coli* BL21 (DE3), except H4, which was expressed with *E. coli* JM109 (DE3). The nucleosome was reconstituted with purified histone octamer and the 145 bp DNA fragment containing Widom 601 sequence^75^ by the salt dialysis method, as described previously.^76^ The reconstituted nucleosomes were further purified by polyacrylamide gel electrophoresis, using Prep Cell apparatus (Bio-Rad).

Thermal stability assays were performed as described previously.^55,77^ The nucleosome containing the 145 base-pair Widom 601 DNA was mixed with 1× Protein Thermal Shift Dye (ThermoFisher), in 18 mM Tris–HCl (pH7.5) buffer containing 100 mM NaCl, 4.5% glycerol, and 0.9 mM DTT. The fluorescence signals of the Protein Thermal Shift Dye were detected with a StepOnePlus Real-Time PCR unit (Applied Biosystems), with a temperature gradient from 26 to 95°C, in steps of 1°C/min. The fluorescence intensity was normalized as follows: [F(T) – F(26°C)/[F(95°C) – F(26°C)]. F(T) indicates the fluorescence intensity at a particular temperature.

### *In Vitro* Pulldown Experiments

Recombinantly expressed and purified proteins were bound to 10 µl Ni-NTA (Qiagen Cat. No. 30210) resin previously equilibrated in pulldown buffer (50 mM Tris pH 8, 150 mM NaCl, 2.5 mM DTT, 0.1% tween-20). Bait proteins were then bound to the prepared resins for 1 hr at RT. For *S. pombe* Hta1 and Pht1 experiments, 10 µg was used. For *S. pombe* Spt6, 25-50 µg was used. For peptide experiments, a vast excess (50 µg) of biotinylated peptide was used. The resins were then washed 4 times with pulldown buffer, then the prey protein was added, incubating for 2 hrs at RT. Beads were then washed 4 times, then proteins were eluted from the beads by boiling in SDS-PAGE running buffer. Proteins were then separated on an SDS-PAGE gel and analyzed by either colloidal blue staining or Western blot as indicated above, using a streptavidin-HRP conjugate (Invitrogen Ca. No. S911) for detection.

### mRNA Sequencing Library Preparation and Analysis

Total RNA was extracted from yeast strains grown to mid-exponential phase (OD_600_ ∼0.5) using a standard hot acid phenol protocol. Libraries were then prepared by polyA-tail enrichment using the NEBNext Ultra II Directional RNA Library Prep Kit for Illumina (NEB Cat. No. E7760L) with the polyA selection module (NEB Cat. No. E7490L). Size selection steps and final library cleanup were performed using the NA clean-up bead solution provided by the VBC core facilities, which is adapted from DeAngelis et al. 1995^78^ using carboxylate-modified Sera-Mag Speed beads (Cytiva). Library quality control was performed using the Advanced Analytical Fragment Analyzer using the HS NGS fragment analysis kit (Agilent Cat. No. DNF-474) as well as qPCR using reagents produced by the Molecular Biology Service in conjunction with commercial DNA standards (Roche Cat. No KK4903). Data was collected on a Thermo Scientific QuantStudio5 instrument.

Prepared libraries were sequenced on either an Illumina NextSeq2000 or NovaSeq S4 at the Vienna BioCenter Core Facilities Next Generation Sequencing Core using paired-end mode. Quality control of the data was performed using FastQC (Babraham Institute). Transcript-level quantification against the *S. pombe ASM294v2 reference genome* available from ENSEMBLv55 was performed using Kallisto.^79^ Differential expression analysis was performed using DESeq2^80^ in R, where all subsequent data manipulation was performed.

### PRO-seq Library Generation and Analysis

We performed a variant of PRO-seq, qPRO-seq, as recently published,^37^ with modifications^38^ to make it compatible with *S. pombe*. *S. pombe* cells were grown in 10 ml YES media to OD_600_ ∼0.4–0.5. Cells were then harvested by centrifugation at 400 x *g* for 5 min at 4 °C. Cells were then resuspended in 10 ml of ice-cold PBS, spun down once more, then resuspended in 10 ml ice-cold yeast permeabilization buffer (0.5% Sarkosyl, 0.5 mM DTT, Roche cOmplete protease inhibitor cocktail, 4U/ml Invitrogen RiboLock RNAse Inhibitor). After 20 min incubation on ice, cells were once again collected by gentle centrifugation, then resuspended in 50 µl storage buffer (10 mM Tris-HCl, pH 8.0, 25% (v/v) glycerol, 5 mM MgCl_2_, 0.1 mM EDTA, 5 mM DTT).

A 2X run-on reaction master mix (40 mM Tris pH 7.7, 64 mM MgCl2, 1 mM DTT, 400 mM KCl, 40 µM Biotin-11-CTP, 40 µM Biotin-11-UTP, 40 µM ATP, 40 µM GTP, 1% sarkosyl) was prepared and preheated to 30 °C. To each aliquot of permeabilized cells, 50 µl of master mix and 1 µl of RNAse inhibitor were added, then mixed by gently pipetting with wide-bore pipet tips before immediately incubating at 30 °C for 5 min. To ensure precise timing, samples were done in batches staggered by 30–60 sec. Using the Norgen RNA Extraction Kit (Cat. No. 37500), 350 µL of RL buffer were added to each sample and then vortexed. 240 µl of absolute ethanol were added, then the entire well-mixed solution was applied to the RNA extraction column. The column was spun at 3500 x *g* for 1 min at 25 °C, then washed once with 400 µl of wash solution A. This step was repeated a second time, then the column was dried by spinning at high speed for 2 min. RNA was eluted twice in 50 µl of DEPC-treated MonoQ water, the two fractions were pooled and then briefly denatured at 65 °C for 30 sec. After snap cooling on ice, 25 µl of ice cold 1 M NaOH were added to each sample, which was then incubated for 10 min on ice. Finally, samples were quenched with 125 µl of cold 1 M Tris-HCl pH 6.8, then cleaned up using a standard ethanol precipitation overnight. Samples were resuspended in a small volume of DEPC water (6 µl).

Meanwhile, 10 µl per sample of Dynabeads MyOne Streptavidin C1 beads (Thermo Fisher Cat. No. 65001) were prepared by washing once with bead preparation buffer (0.1 M NaOH, 50 mM NaCl), then once in bead binding buffer (10 mM Tris-HCl, pH 7.4, 300 mM NaCl, 0.1% (v/v) Triton X-100, 1 mM EDTA, RNAse inhibitors), then resuspended in 25 µl bead binding buffer.

3’ RNA adapter ligation was performed by adding 1 µl 10 µM oligo VRA3 to the resuspended RNA, boiling at 65 °C for 30 sec, then snap cooling on ice. Ligation reactions were then prepared with T4 RNA Ligase using the provided instructions. After incubating reactions at 25 °C for 1 hr, ligated RNA was bound to the 25 µl previously prepared streptavidin beads by adding 55 µl bead binding buffer and the entire ligation reaction. Samples were incubated for 20 min at 25 °C, then washed once with 500 µl high salt wash buffer (50 mM Tris-HCl, pH 7.4, 2 M NaCl, 0.5% (v/v) Triton X-100, 1 mM EDTA, 2 µl/10 ml Ribolock RNAse inhibitor), transferring beads to a clean tube. Beads were then washed once more in the new tube with low salt wash buffer (5 mM Tris-HCl, pH 7.4, 0.1% (v/v) Triton X-100, 1 mM EDTA, 2 µl/10 ml Ribolock RNAse inhibitor), then resuspended in 20 µl T4 PNK reaction mix (1X T4 PNK Buffer, 1 mM ATP, 1 µl T4 PNK, 1 µl RiboLock RNAse inhibitor). Samples were then incubate at 37 °C for 30 min, then the beads were collected by magnet and resuspended in 20 µl ThermoPol reaction mix (1X ThermoPol buffer, 1 µl RppH, 1 µl RiboLock RNAse inhibitor), then incubated at 37 °C for 1 hr. Beads were once again collected by magnet, then the 5’ RNA adaptor (REV5) was ligated using T4 RNA ligase, setting up the reaction using the provided manual and incubating at 25 °C for 1 hr. Beads were then washed once with high salt wash buffer, transferring to a clean tube, then once with low salt buffer, before being resuspended in 300 µl of TRIzol reagent (Ambion). Samples were vortexed for 20 sec, then incubated on ice for 3 min before adding 60 µl of chloroform, vortexing for 15 sec, then incubating on ice for a further 3 min. Samples were then spun at 20,000 x *g* for 8 min at 4 °C, then the upper ∼180 µl aqueous phase was transferred to a clean tube and precipitated using a standard ethanol precipitation.

Finally, pellets were resuspended in 13.5 µl reverse transcription resuspension mix (8.5 µl DEPC water, 4 µl 10 µM primer RP1, 1 µl 10 mM dNTPs), denaturing at 65 °C for 5 min, snap cooling on ice. Reverse transcription was then performed using Maxima H Minus RT (Thermo Fisher Cat. No. EP0753) by adding 6.5 µl RT master mix (4 µl 5X RT buffer, 1 µl DEPC water, 0.5 µl RiboLick RNAse inhibitor, 1 µl Maxima H Minus RT), then cycling as follows: 50 °C for 30 min, 65 °C for 15 min, 85 °C for 5 min. 2.5 µl of indexed RPI-n primer (10 µM) were then added to each RT reaction, then PCR was performed as a 100 µl reaction using the Q5 polymerase (NEB) with the included high GC content enhancer. Samples were then PCR-amplified to the beginning of the logarithmic phase, as determined using a small-scale qPCR reaction run in parallel (typically less than 20 cycles).

Libraries were sequenced as described previously at the Vienna BioCenter Core Facilities Next Generation Sequencing Core as previously. The data analysis was performed by adapting available pipelines.^37^ Briefly, we created a Nextflow container (https://github.com/Gregor-Mendel-Institute/PROalign) to perform all QC, alignment, and read processing steps, generating strand-specific bigwig files that were subsequently analyzed in R.

### ChIP-seq and -qPCR

*S. pombe* strains (100 ml) were grown to mid-exponential phase (OD_600_ ∼0.4-0.5) in YES, fixed with 1% formaldehyde for 10 min, quenched with 125 mM glycine for 10 min, then collected by centrifugation. Cells were washed once with ice-cold PBS, then resuspended in lysis buffer (50 mM HEPES-KOH, pH 7.5, 140 mM NaCl, 1 mM EDTA, 0.1% sodium deoxycholate, 1% Triton X-100, 1X Roche Complete Protease Inhibitors), and then lysed using acid-washed glass beads in a Precellys for 4×20 sec maximum speed, 1 min on ice between rounds. Lysate was then separated from the beads and then chromatin was sheared to a predominant final DNA length of ∼150-200 bp in a Covaris E220 ultrasonicator, and then clarified of unlysed cells and debris by centrifugation twice at 16,000 × *g* for 10 min. Prepared chromatin was then frozen at −70 °C pending use.

Chromatin aliquots were thawed on ice, then normalized based on starting culture density using additional lysis buffer for dilution. 10% of normalized and diluted chromatin were reserved as input controls. Anti-H2A.Z (Pht1) or anti-Myc 9E10 antibody (1µg) was then added to each culture and samples were incubated overnight at 4 °C with inversion. 25 µl of Protein A Dynabeads (Thermo Fisher Cat. No. 10001D) per sample were prepared by washing twice with PBS with 0.1% tween-20, once with lysis buffer, then resuspending in 100 µl of lysis buffer. Prepared beads were then combined with overnight chromatin samples, and incubated for 4 hrs at 4 °C with inversion. Beads were then collected using the magnet and washed twice with lysis buffer, twice with high salt lysis buffer (50 mM HEPES-KOH, pH 7.5, 500 mM NaCl, 1 mM EDTA, 0.1% sodium deoxycholate, 1% Triton X-100), twice using lithium wash buffer (10 mM Tris-HCl, pH 8, 250 mM LiCl, 1 mM EDTA, 0.5% NP-40, 0.5% sodium deoxycholate), and once with TE. Beads were then resuspended in 100 µl of elution buffer (50 mM Tris-HCl, pH 8, 10 mM EDTA, 0.8% SDS) and transferred to PCR strips. Samples were then boiled for 10 min at 95 °C, then 65 °C overnight in a thermocycler. The next morning, proteinase K was added (final concentration 0.2 mg/mL) and incubated at 55 °C for 2 h. Samples were then cleaned up using a commercial kit (Zymo Research Cat. No. D5201). DNA concentration was measured by Nanodrop spectrophotometer. To verify the H2A.Z antibody, ChIP-qPCR was set a 100-fold dilution of samples was used in a 15 µl reaction using a 2X SYBR-green master mix provided by the IMP Molecular Biology Service. For the *ura4*^+^ locus, primers ZH_1908 (5’-GCAGGTCACAGTCATGAAGCCA-3’) and ZH_1909 (5’-GGATTTTTCATCCCCTCAGCTCTAGC-3’) were used.

For library preparations, all samples, including input, were processed using the NEBNext Ultra II library preparation kit for Illumina (NEB Cat. No. E7645L) using custom synthesized barcoded sequencing adapters provided by the Vienna BioCenter Core Facilities Next Generation Sequencing Facility. Following quality control by Agilent Fragment Analyzer and quantification by RT-qPCR using a Kapa Library Quantification Kit (Roche Cat. No. KK4903), libraries were sequenced at the Next Generation Sequencing Facility on an Illumina NovaSeq X. Alignments, quality control, and initial processing were performed using the nf-core/chipseq pipeline (https://github.com/nf-core/chipseq) with default parameters. Input-normalized bigWig files were subsequently generated using Deeptools (v3.3.1).^81^

### Quantification and Statistical Analysis

Unless otherwise noted, all statistical analyses were performed in R (v4.4.2) using the RStudio IDE (2024.12.0+467) on Darwin 24.3 (64-bit) under macOS 15.3.1. All statistical tests used are stated in corresponding figure legends. Quantification of sequence distances was performed using CLC Main Workbench 10 (Qiagen). AlphaFold2 Multimer^43^ analyses were performed using ColabFold^82^ on the CLIP cluster (https://clip.science).

### Data Availability

Datasets from phylogenetic analyses are provided as supplementary data files. PRO-seq pipeline and scripts used for data analysis are available as a container on Github (https://github.com/Gregor-Mendel-Institute/PROalign), and all other scripts used in analysis are available upon request to the lead contact. NGS data are deposited in GEO under accession numbers: PRO-seq (GSE265749), mRNA-seq (GSE265775), and ChIP-seq (GSE279717, GSE280242).

## Acknowledgements

We thank the entire Berger group and Antoine Hocher, for their insightful, considerate, and helpful discussions, as well as Carol Featherstone from Life Science Editors and Matt Watson for editorial assistance during the preparation of the manuscript. We especially thank Zdravko Lorković, Elin Axelsson-Ekker, and Chung Hyun Cho for their invaluable assistance preparing antibodies, biochemical assays, and bioinformatic analyses. We further thank Hitoshi Kurumizaka, Takumi Oishi, Rina Hirano, and Suguru Hatazawa for their assistance providing materials to prepare nucleosomes. We also thank John T. Lis for advice with qPRO-seq. We thank the Molecular Biology Service and Media Kitchen for a constant supply of plates, basic reagents, and cloning and sequencing services, as well as the Peptide Synthesis Service. Additionally, we thank the Vienna BioCenter Core Facilities, in particular the Next Generation Sequencing facility for their advice and swift handling of all our requests and the Protein Technologies facility for their expertise, helpful discussions, and recombinant protein production.

## Funding

Gregor Mendel Institute, Austrian Academy of Sciences Core Funding (FB)

Österreichischer Wissenschaftsfonds FWF TAI304 (FB)

Österreichischer Wissenschaftsfonds FWF ESP213 (ZHH)

EMBO Postdoctoral Fellowship ALTF169-2020 (ZHH)

United Kingdom Research and Innovation’s Medical Research Council MC_A658_5TY40 (TW)

JSPS KAKENHI JP24K02002, JP24H01351 (AO)

JST PRESTO JPMJPR20K3 (AO)

JST CREST JPMJCR20S6 (AO)

## Author contributions

Conceptualization: ZHH, FB, and TW; Methodology: ZHH, KS, JYK, AO, JL, and TW; Validation: ZHH, KS, JYK, and AO; Formal Analysis: ZHH, KS, and TW; Investigation: ZHH, KS, JYK, AO, JL, and TW; Data Curation: ZHH, KS, JYK, and TW; Writing – original draft: ZHH, TW, and FB; Visualization: ZHH, KS, and TW; Supervision: FB, TW; Project Administration: ZHH; Funding Acquisition: ZHH, FB, AO, and TW.

## Competing Interest

Z.H.H. and F.B. are inventors on a patent submitted by the Gregor Mendel Institute at the Austrian Patent Office (A50392/2024) pertaining to the modification of transcription and fitness by engineering H2A.Z L2.

## Materials and Correspondence

For material request (e.g. strains), please contact the lead.

---

**Extended Data Figure 1.**
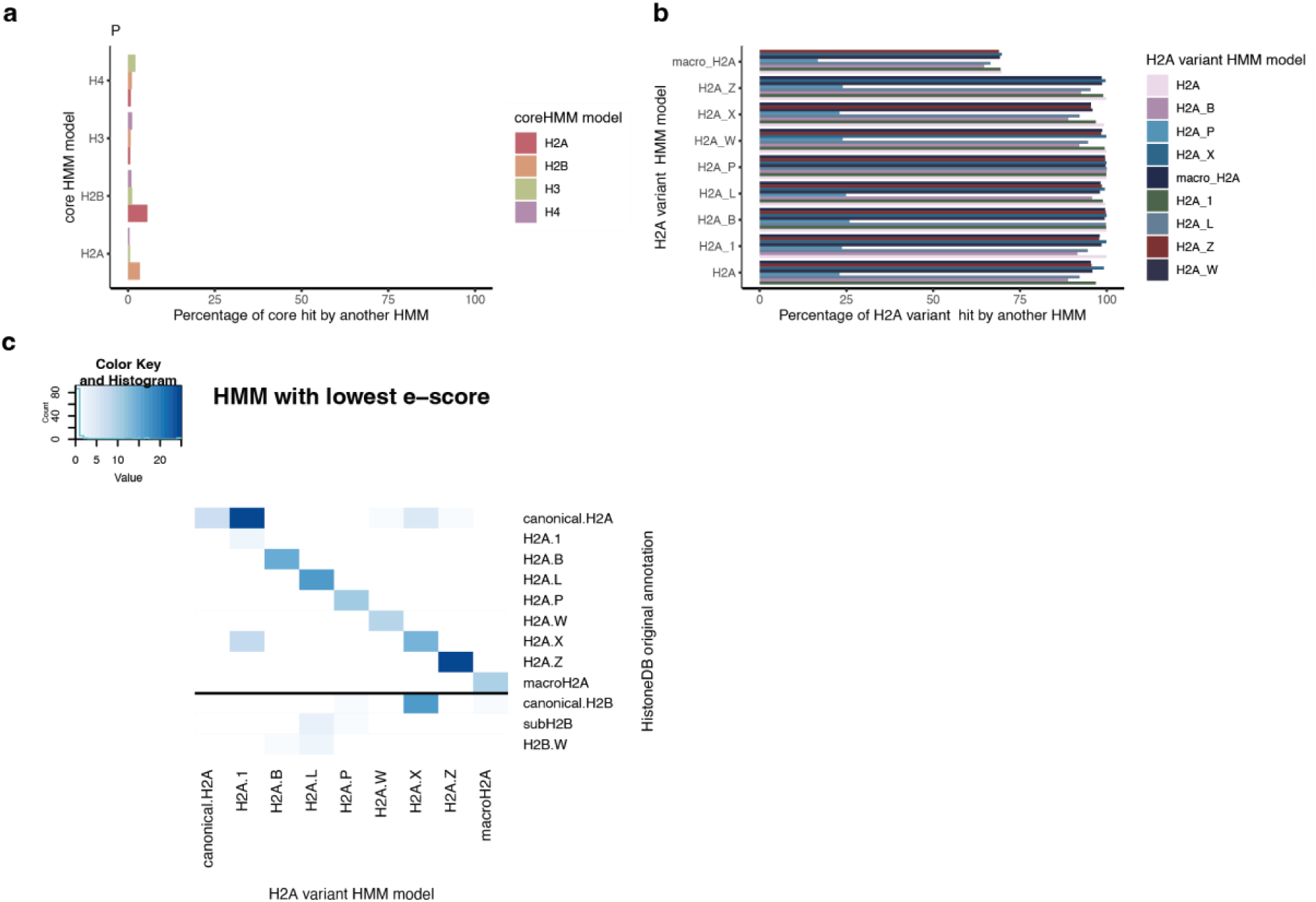
Validation of HMM Models Used to Annotate Histone Variants. **a,** Specificity of core histone HMMs queried against histones in EukProt. **b,** Specificity of H2A variant HMM models queried against histones in EukProt. **c,** H2A variant HMMs do not reliably identify the right H2A variant, even if assigning variant identify based on the lowest E-value.

**Extended Data Figure 2.**
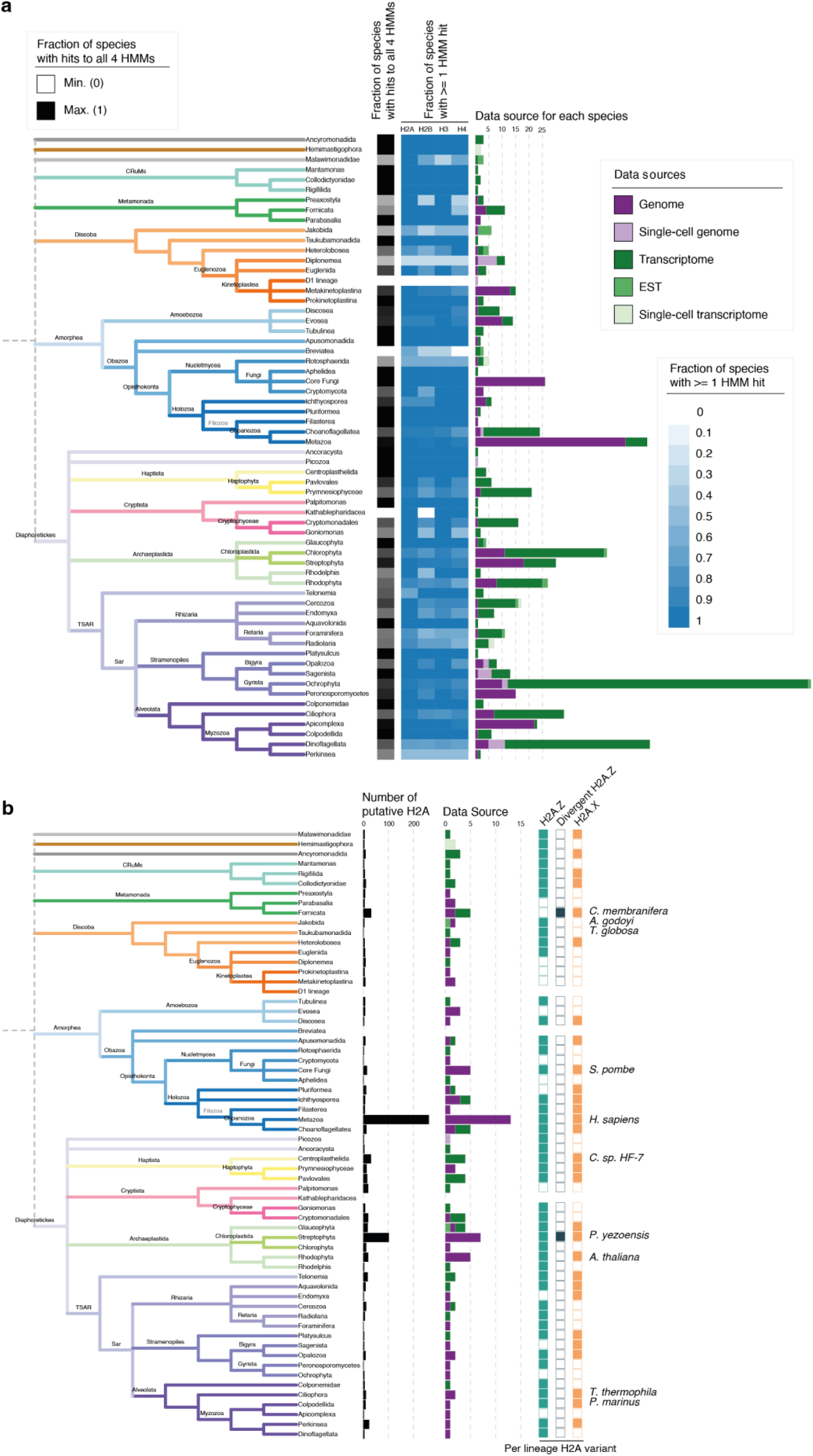
Overview of H2A Dataset. **a,** Overview of histones identified by different core histone HMMs in the context of eukaryotic phylogeny. Number of representative genomes and their data source are also indicated. **b,** Number of H2As and presence/absence of specific variants (H2A.Z, H2A.X, divergent H2A.Z, as mentioned in the main text) across eukaryotic phylogeny. Numbers reflect the distribution in the final filtered dataset. The cladogram of major eukaryotic lineages and information on data sources are taken from EukProt. H2A.Z from the list of species in the far-right column were chosen for further investigation.

**Extended Data Figure 3.**
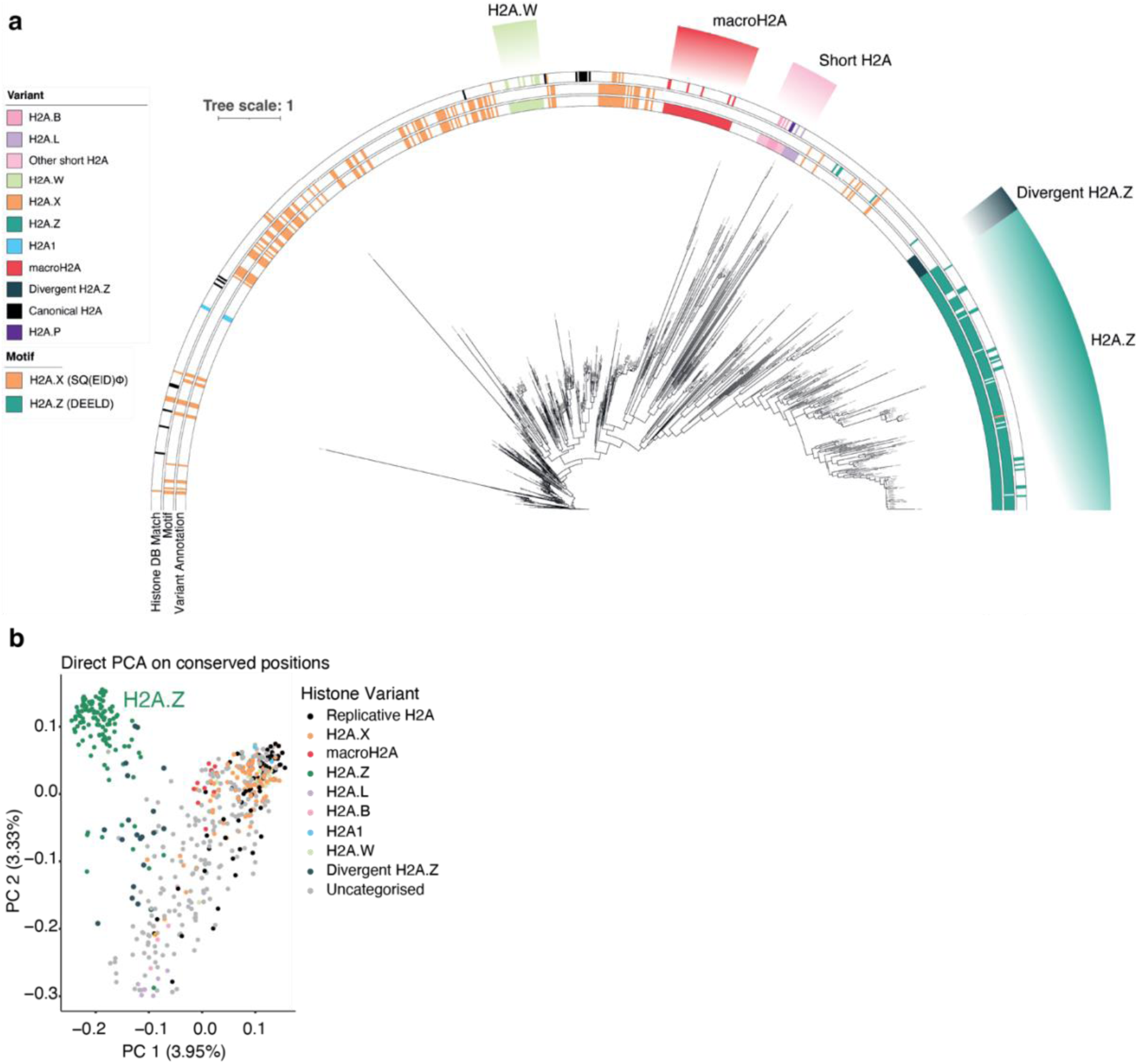
H2A/Z Molecular Phylogeny. **a**, Unrooted protein phylogeny of H2A (see Methods for reconstruction details). H2A.Z, as defined by the presence of a ‘DEELD’ motif within its docking domain, are colored in green. ‘Divergent’ H2A.Zs branch as a sister to the major H2A.Z clade and carry at least one substitution in the DEELD motif. **b**, Principal component analysis (PCA) on H2A sequences displayed in **a**, annotated by variant identity.

**Extended Data Figure 4.**
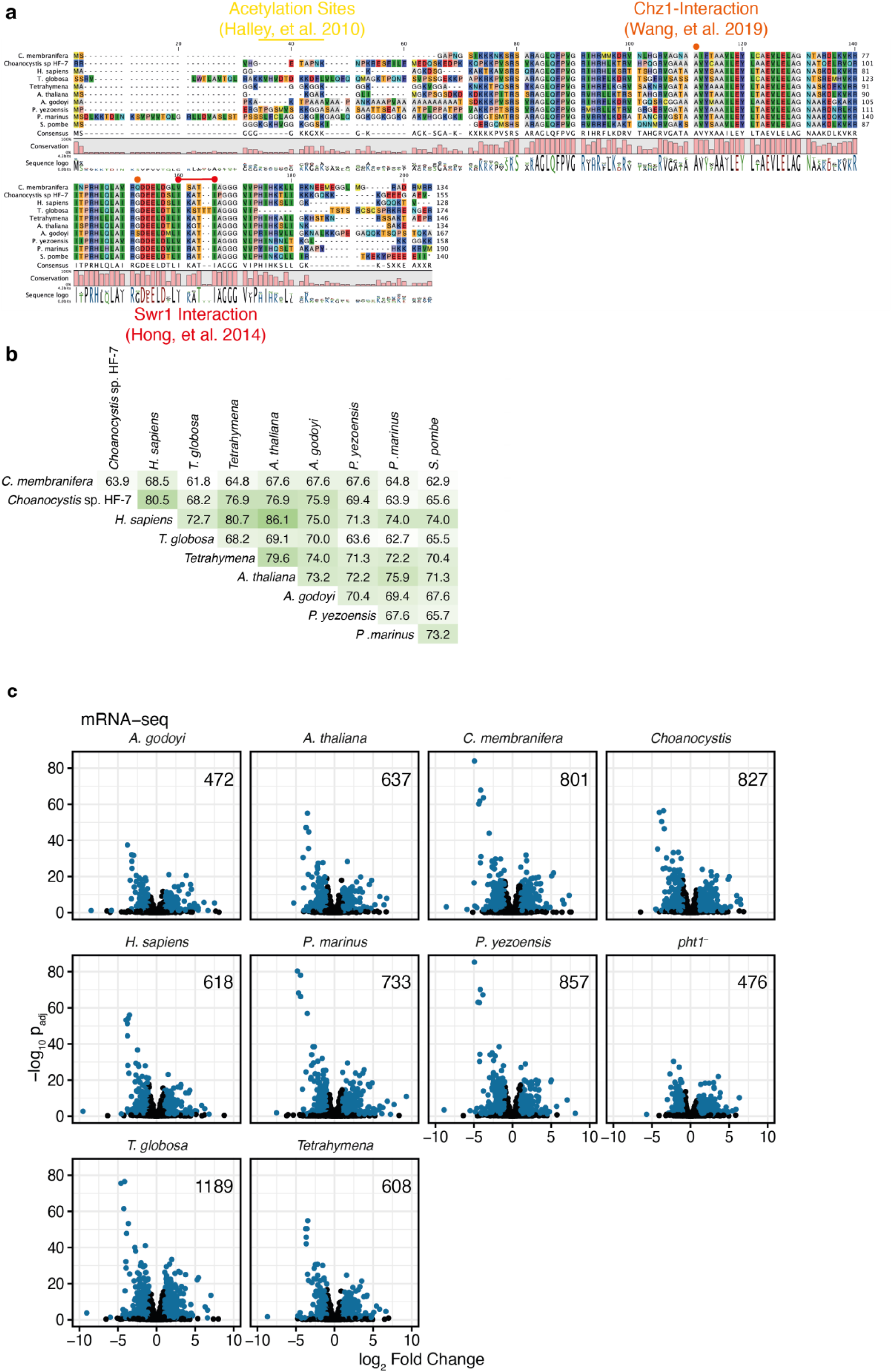
Additional Analysis of Pht1 Replacement Line Sequence and Transcriptomes. **a,** Alignment of representative H2A.Z sequences taken from our phylogeny, with key residues and relevant literature noted. **b,** Pairwise percent identity of representative H2A.Zs, excluding N- and C-terminal tail regions. **c,** Differential expression analysis of mRNA-seq data for each species’s H2A.Z replacement line against WT *S. pombe*. Number of differentially expressed genes (*p_adj_*<0.1, out of 5134 coding genes) is indicated for each panel.

**Extended Data Figure 5.**
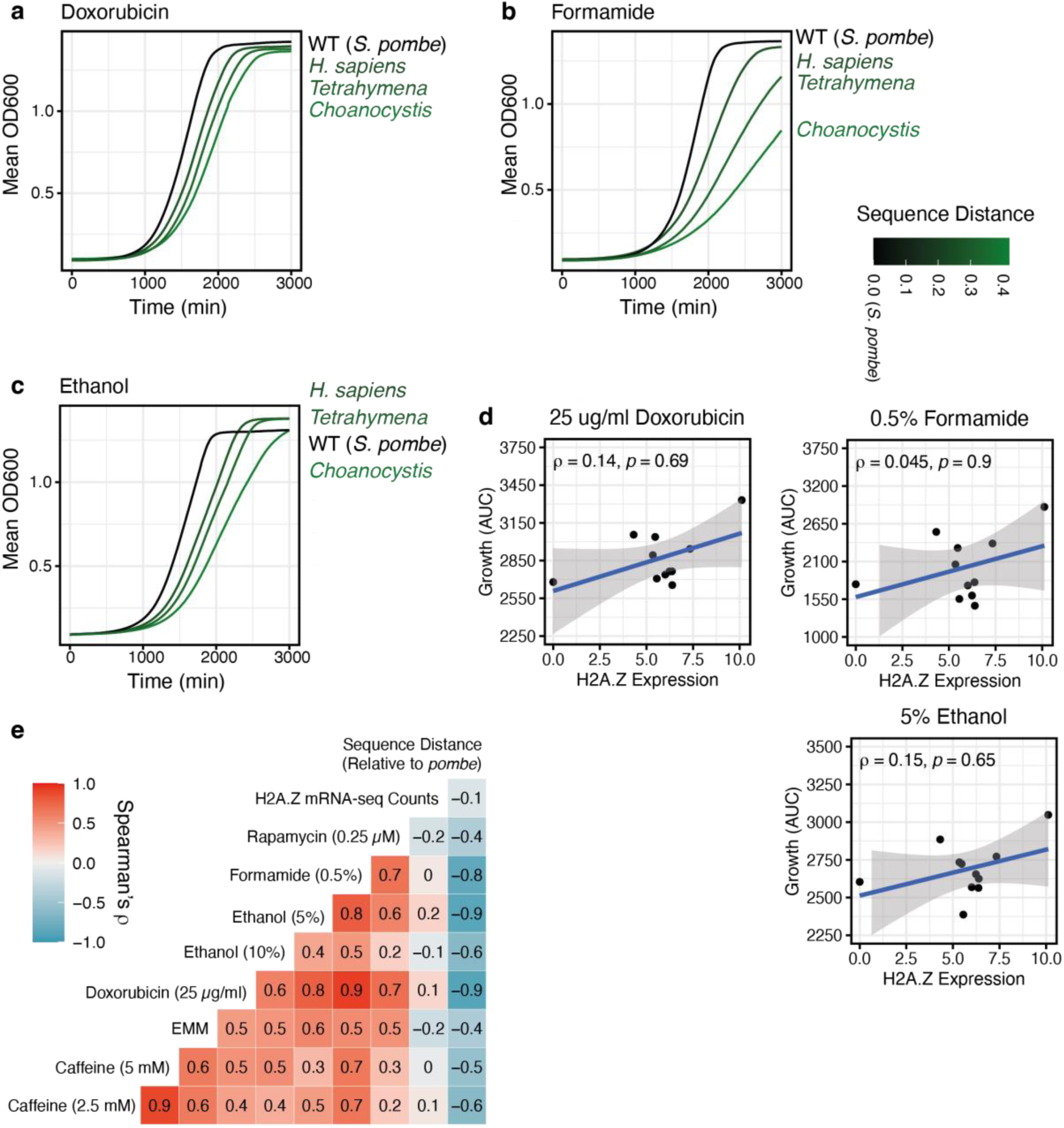
Additional Phenotypic Data and Analysis for Full-Length Pht1 Replacement Lines. **a–c,** Example growth curves related to Fig. 1c for selected phenotypes and strains. Curves are the mean of 4 biological replicates and are coloured according to the evolutionary distance of the H2A.Z from a given species. **d,** Example Spearman’s correlation analysis of sequence distance relative to WT *S. pombe* Pht1 relative to Pht1 mRNA levels from Fig. 1b for each of the 9 full-length H2A.Z replacement lines. **e,** Further Spearman’s correlations for the phenotypes of H2A.Z replacement lines from Fig. 2b with themselves, as well as either their sequence distance from WT *S. pombe* or their expression.

**Extended Data Figure 6.**
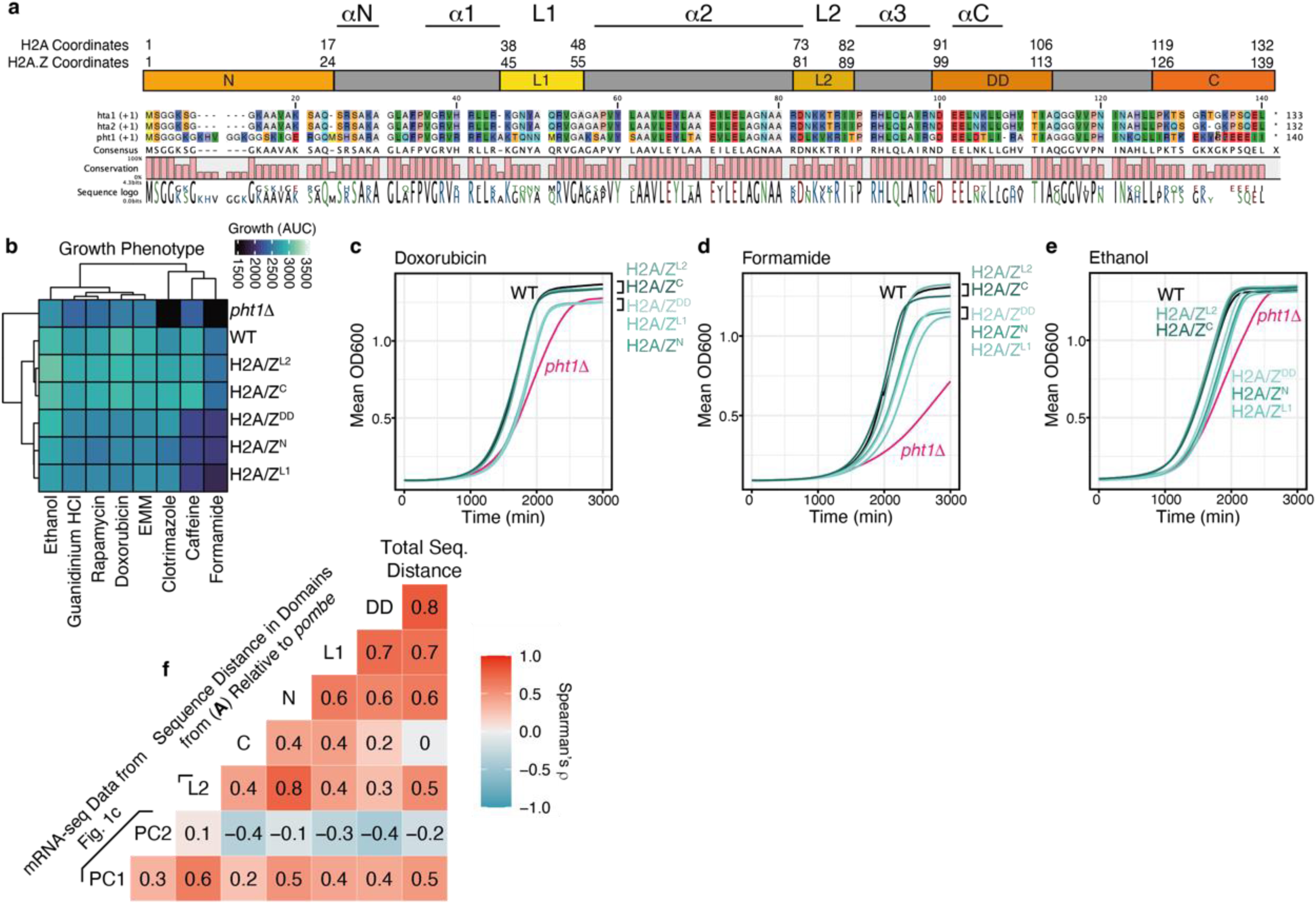
Domain Analysis of Pht1. **a,** Alignment of *S. pombe* H2A and H2A.Z identifying the five regions. **b**, Mean growth (AUC) for four biological replicates in each indicated condition, which are as in Fig. 1B for S. pombe strains encoding H2A/H2A.Z chimeras according to the regions defined in **a**. **c–e,** Example growth curves related to the heatmap from **b**. Plotted is the mean of 4 biological replicates for the indicated strain, and the conditions are as in Fig. 1c. **d,** Additional Spearman’s correlation analyses of mRNA-seq data from Fig. 1b against the sequence distance for each complemented H2A.Z in the indicated regions from **a**.

**Extended Data Figure 7.**
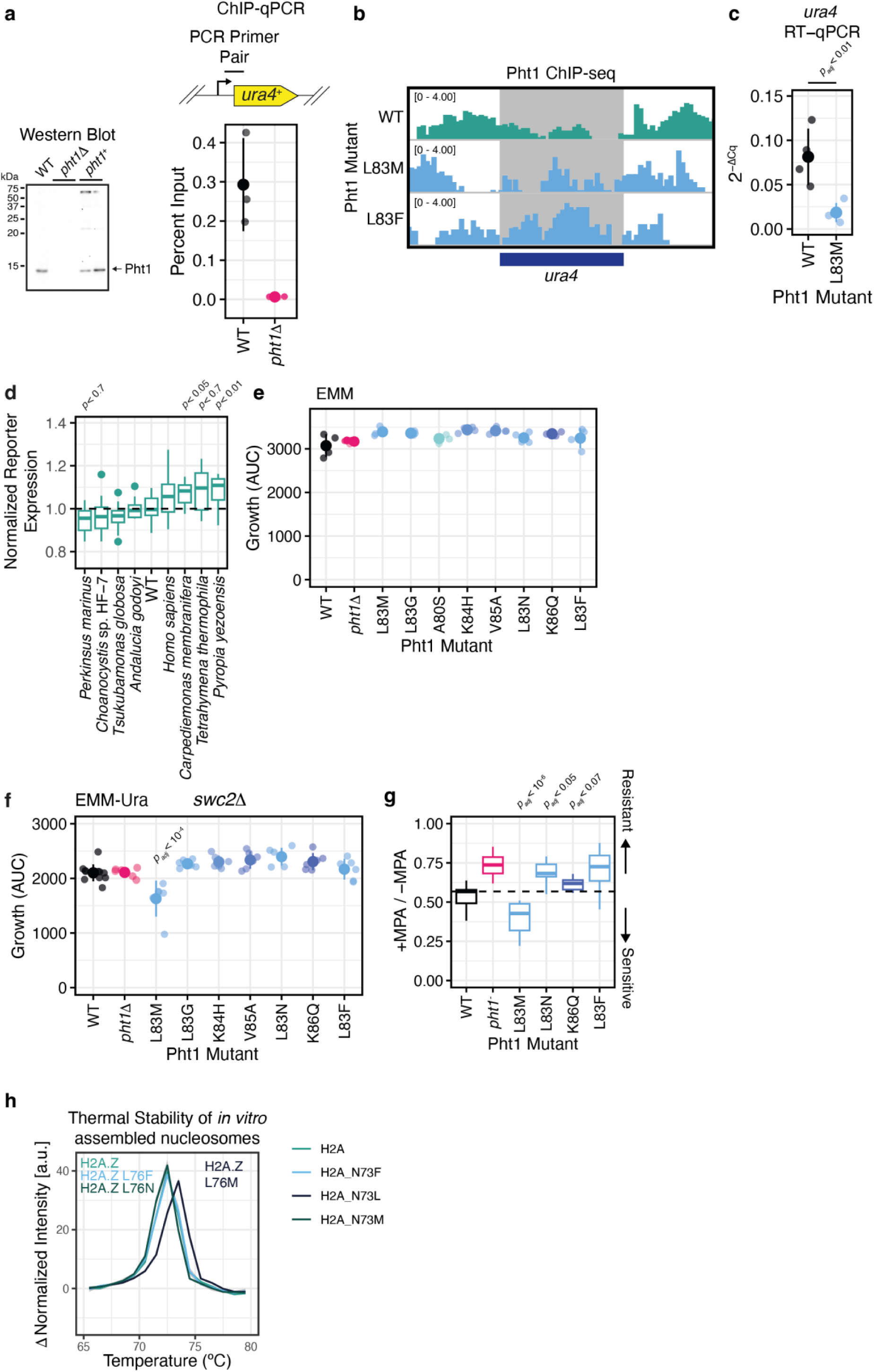
Controls Pertaining to Transcription Measurements. **a,** ChIP-qPCR data for *ura4* reporter in either WT and *pht1*Δ cells. Plotted are data from three independent ChIP experiments and are the average of 4 technical replicates. Accompanying western blot to validate the selectivity of the Pht1 antibody, performed against acid extracts of WT, *pht1*Δ, and *pht1*^+^, whereby *pht1* was knocked back into the *pht1*Δ background. **b,** Pht1 ChIP-seq profiles for WT, Pht1 L83M, and L83F for the *ura4* reporter locus. **c,** RT-qPCR analysis for indicated strains against *ura4*^+^ mRNA. Quantification performed using *S. pombe act1* as a reference. **d,** growth of *ura4* reporter strains with the Pht1 L2 replaced at the endogenous locus for the L2 for the indicated species. Plotted are at least 10 biological replicates, and *p-*values are calculated with a Wilcoxon test relative to WT *S. pombe*. **e,** Growth of H2A.Z point mutant strains in complete growth medium (EMM). Plotted are four biological replicates, with their mean and standard error. **f,** *ura4* reporter assay for L2 point mutants from Fig. 4C in a *swc2*Δ background. Plotted are at least 5 biological replicates with their mean and standard error. **g,** Growth of indicated *S. pombe* Pht1 mutants in the presence of 12.5 µM mycophenolic acid (MPA) expressed as the ratio of their paired growth in the absence of the drug. Data are nine biological replicates and the experiment was repeated three times with similar results. Boxplots represent the median bounded by quartiles. Adjusted *p*-values (*p_adj_*) are calculated using a two-way ANOVA followed by a post-hoc Tukey test. **h,** Thermal stability assays^55^ for H2A.Z L2 neomorphs. Plotted is the mean of three independent experiments bounded by the standard deviation for the derivative of fluorescence intensity.

**Extended Data Figure 8.**
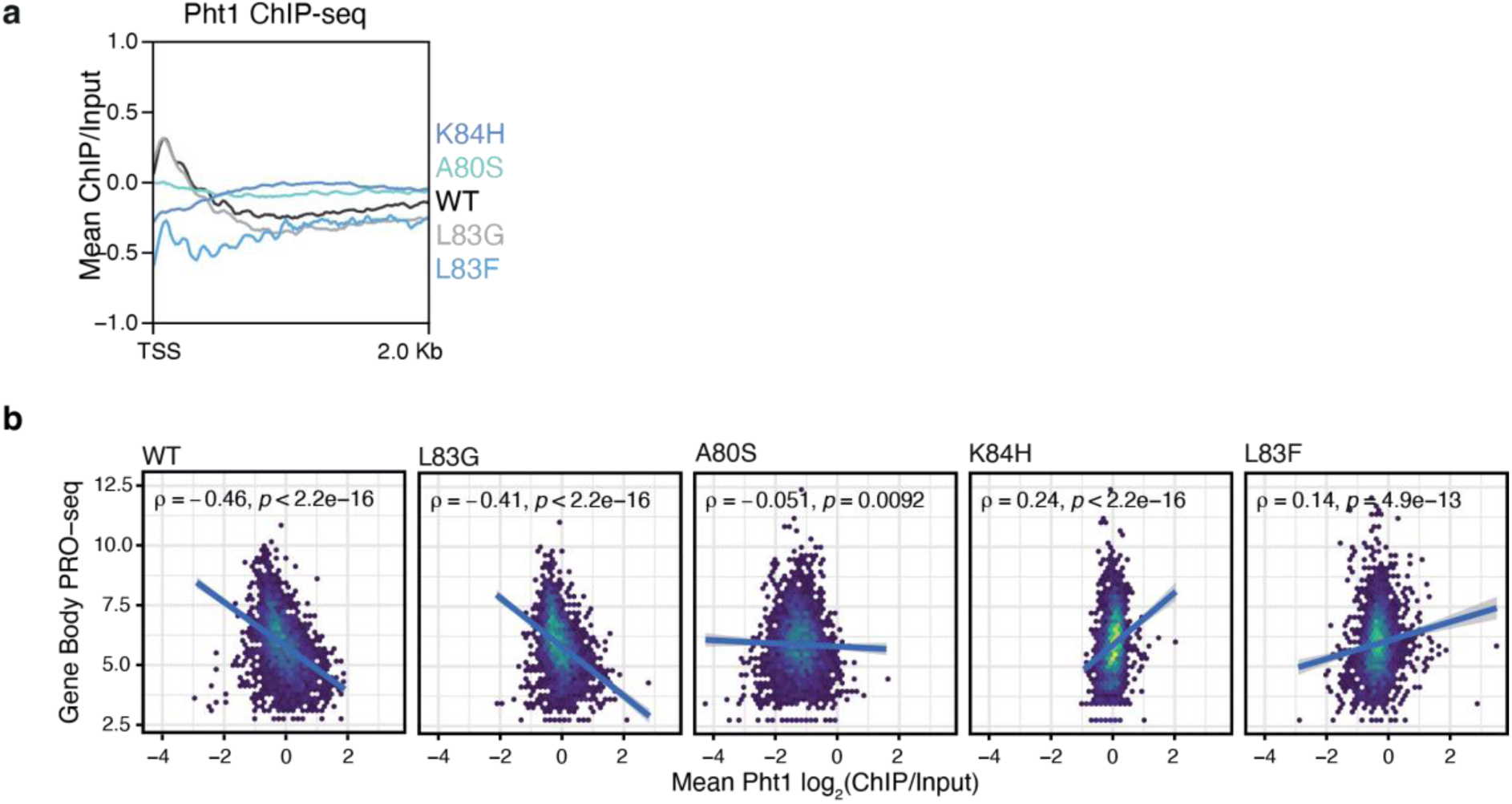
Genomic Profiles of Pht1 L2 Neomorphs. **a,** Example ChIP-seq profiles for Pht1 L2 neomorphs and WT across all transcribed genes referenced to the transcription start site (TSS). Profile is the mean of log_2_-transformed, input normalized signal. **b,** Spearman’s correlation analysis for Pht1 levels genome-wide against transcription as measured by gene-body nascent transcript counts from qPRO-seq for both WT and Pht1 L2 neomorphs.

**Extended Data Figure 9.**
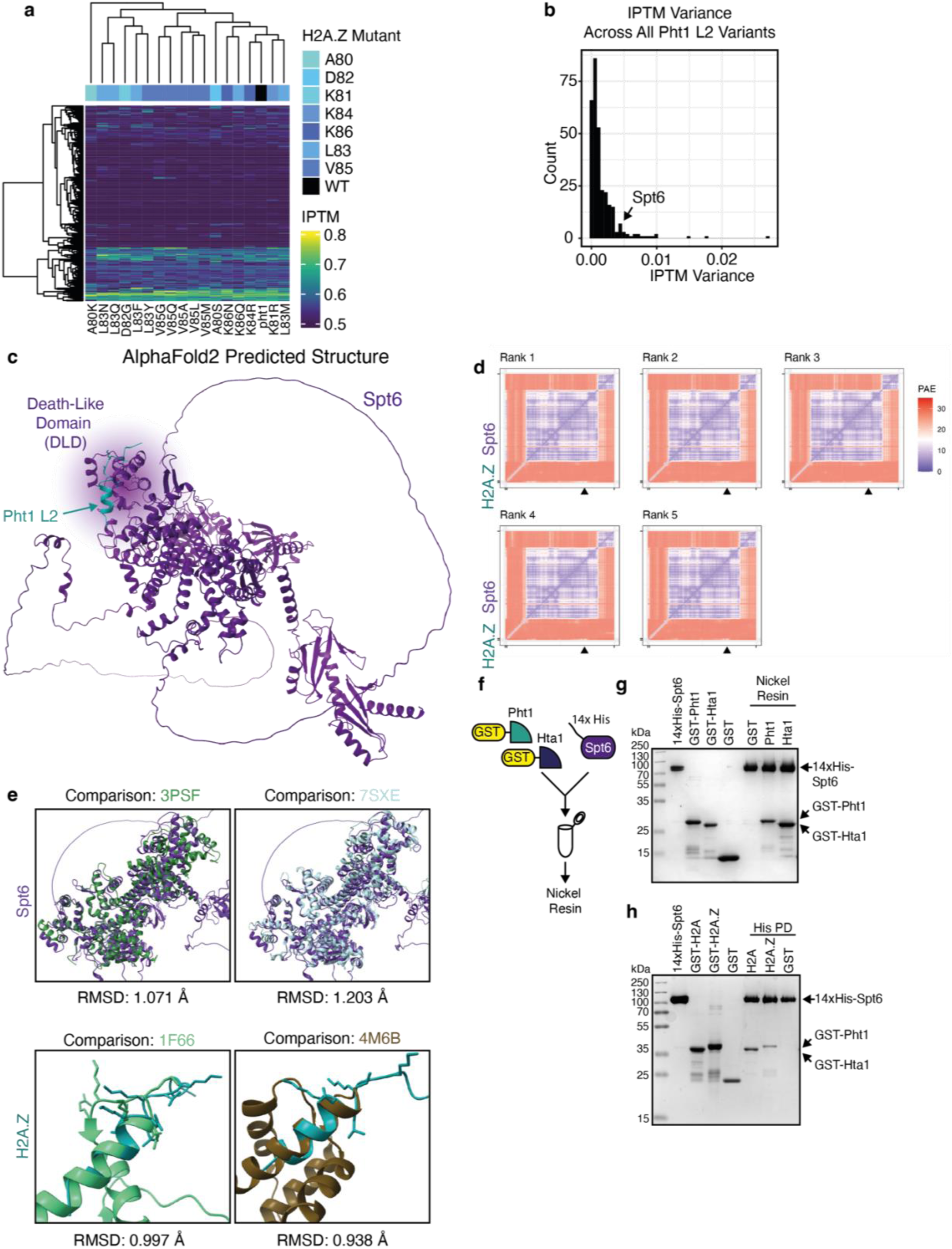
*In Vitro* and *In Silico* Evidence for L2 Neomorph Mechanisms. **a,** AlphaFold2 predicted interaction scores (IPTM) for the top-500 predicted H2A.Z-interacting proteins in *S. pombe* across all tested H2A.Z L2 mutants, as well as H2A and WT H2A.Z. **b,** IPTM variance calculations based on the data from (a), with Spt6 highlighted with an arrow. **c,** Example AlphaFold2-predicted structure for the interaction between Pht1 and Spt6. **d,** AlphaFold2 predicted error alignment (PAE) for the five replicate models of WT *S. pombe* Pht1 against *S. pombe* Spt6. **e,** Comparison of AlphaFold2 predicted *S. pombe* Spt6 structure to either the crystal structure of *S. cerevisiae* (PDB 3PSF) or Cryo-EM structure of *Pichia pastoris* Spt6 (7XSE), as well as predicted Pht1 L2 against the structure of either *H. sapiens* H2A.Z.1 (PDB 1F66) or *S. cerevisiae* Htz1 (PDB 4M6B). **f,** experimental scheme for *in vitro* pulldown experiments for recombinantly expressed and purified 14xHis-Spt6(282– 1159) against Pht1 (H2A.Z). In this assay, Hta1 (H2A) serves as a positive control, and GST a negative control. **g**, SDS-PAGE analysis of *in vitro* pulldown under physiological salt (150 mM NaCl) and **h** under stringent (300 mM NaCl) washing conditions. Detection was performed with colloidal coomasie staining, and was repeated three times with similar results.

**Extended Data Figure 10.**
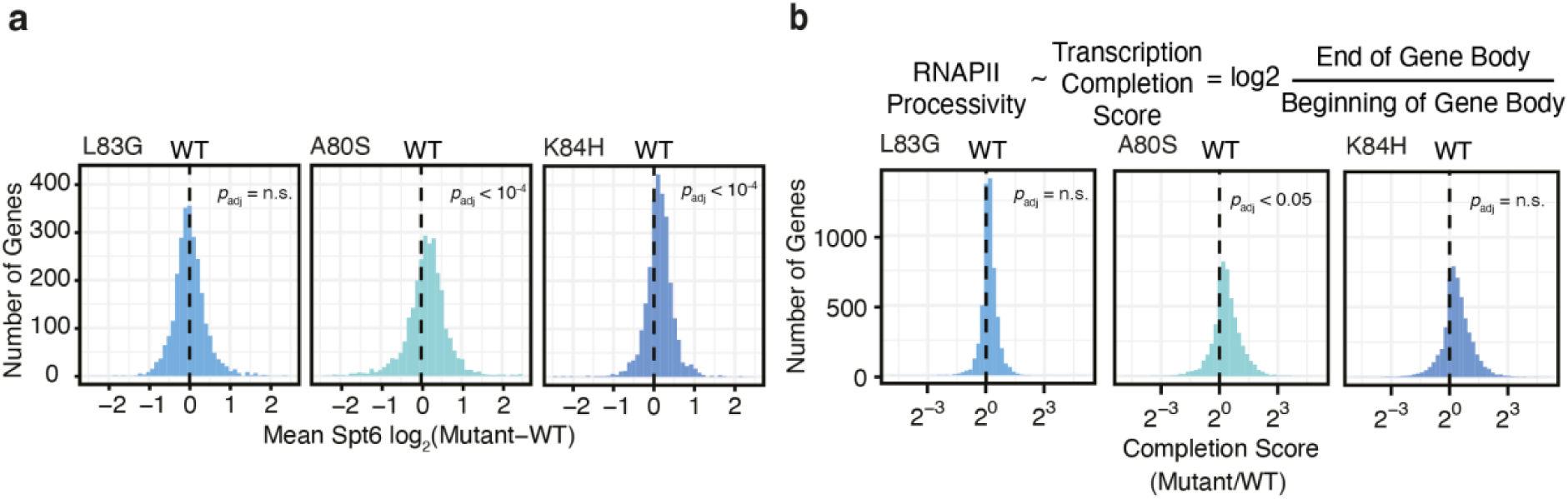
Genomic Analysis for Additional Pht1 L2 Neomorphs. **a,** Analysis of Spt6 levels in indicated Pht1 L2 mutants relative to WT as in Fig. 4g **& h**. Adjusted *p-*values are calculated as a post-hoc Tukey test of a 2-way ANOVA. **b,** Transcription completion score analysis as in Fig. 4i, but for additional Pht1 L2 neomorphs. Adjusted *p*-values calculated as in **c**.

**Extended Data Figure 11.**
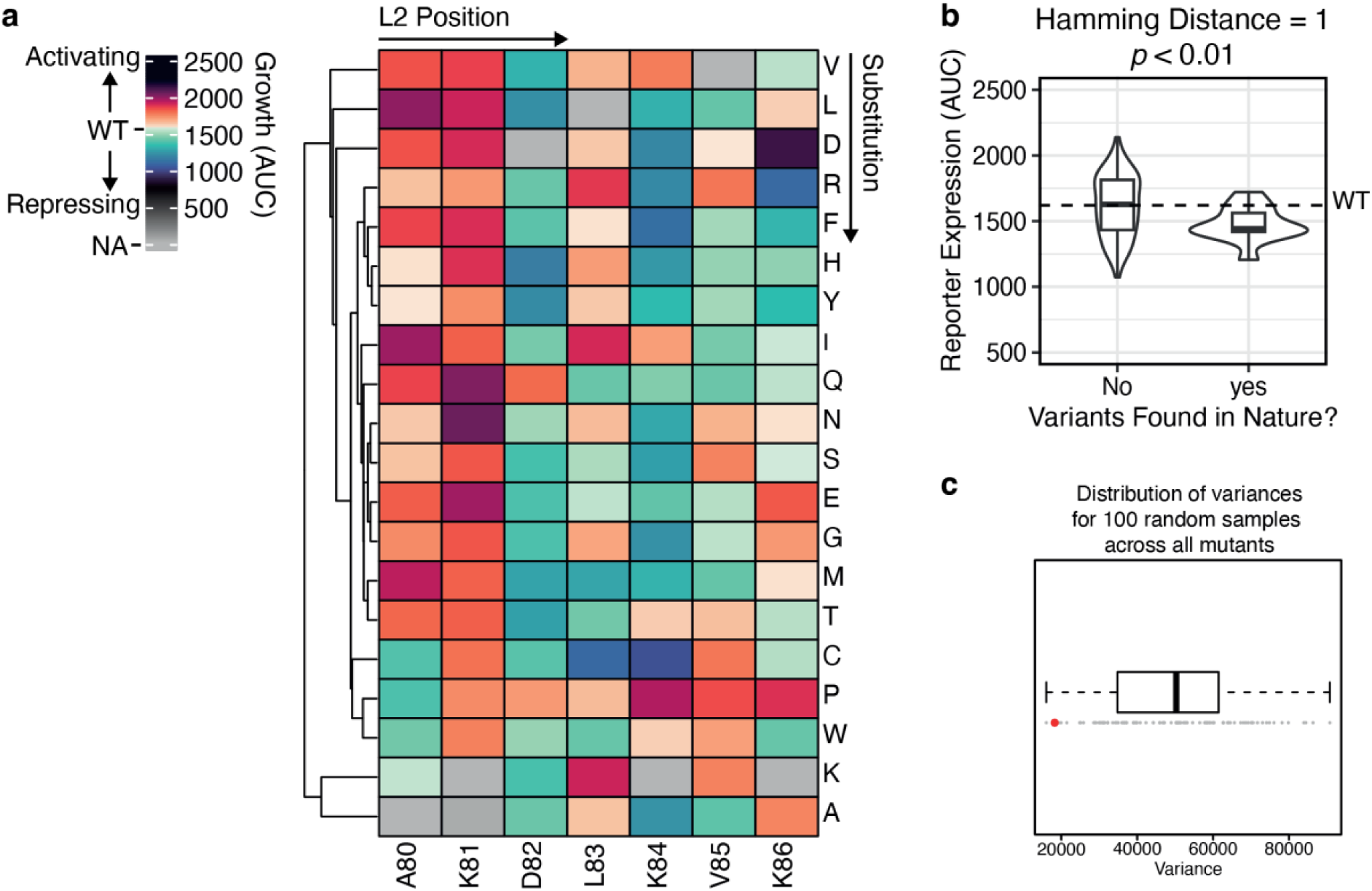
Deep Scanning Mutagenesis of Pht1 L2. **a,** heatmap depicting each possible substitution for each position within the 7-residue L2 core motif coloured according to transcription of the *ura4* reporter relative to WT. Data are the mean of 8 biological replicates. **b–c,** analysis of L2 variants according to whether or not they are present in H2A.Z sequence database at a Hamming distance of 1 relative to WT *S. pombe*. **c**, control simulation for the relatively small number of variants found in nature. 12 variants were randomly selected from the total 133 possible variants 100 times, and their variance calculated as compared to the actual experimentally observed data (red dot). Only one simulation had a lower variance than that observed in the actual data.

## Supplementary Data Files and Tables

**Data Tables S1. (separate file)** Pan-H2A dataset annotations, related to Supplementary Data File S1.

**Data Table S2. (separate file)** AlphaFold2 multimer IPTM vs. PTM predictions for *S. pombe* Pht1 L2 neomorphs against *S. pombe* nuclear proteins.

**Data Table S3. (separate file)** AlphaFold2 multimer IPTM vs. PTM predictions for H2A.Z L2 mutants against *S. pombe* nuclear proteins.

**Data Table S4. (separate file)** Analysis of AlphaFold2 multimer predictions from Data Table S3, with Spt6 highlighted.

**Data Table S5**. **(separate file)** *Schizosaccharomyces pombe* strains used in this study.

**Data S1. (separate file)**

Pan-H2A sequence database used in this study.

**Data S2. (separate file)**

Representative H2A.Z sequences used in this study.

## Notes

### Summary of Updates

We have re-organized all main and supplementary figures, providing substantial new data to support our conclusions. We have also reworked the introduction and discussion.

## References

1 Malik, H. S. & Henikoff, S. Phylogenomics of the nucleosome. Nat Struct Biol 10, 882–891 (2003). 10.1038/nsb996

2 Macadangdang, B. R. et al. Evolution of histone 2A for chromatin compaction in eukaryotes. Elife 3 (2014). 10.7554/eLife.02792

3 Rogakou, E. P., Pilch, D. R., Orr, A. H., Ivanova, V. S. & Bonner, W. M. DNA double-stranded breaks induce histone H2AX phosphorylation on serine 139. J Biol Chem 273, 5858–5868 (1998). 10.1074/jbc.273.10.5858

4 Venkatesh, S. & Workman, J. L. Histone exchange, chromatin structure and the regulation of transcription. Nat Rev Mol Cell Biol 16, 178–189 (2015). 10.1038/nrm3941

5 Kujirai, T. & Kurumizaka, H. Transcription through the nucleosome. Curr Opin Struct Biol 61, 42–49 (2020). 10.1016/j.sbi.2019.10.007

6 Kornberg, R. D. & Lorch, Y. Primary Role of the Nucleosome. Mol Cell 79, 371–375 (2020). 10.1016/j.molcel.2020.07.020

7 Valencia, A. M. & Kadoch, C. Chromatin regulatory mechanisms and therapeutic opportunities in cancer. Nat Cell Biol 21, 152–161 (2019). 10.1038/s41556-018-0258-1

8 Berta, D. G. et al. Deficient H2A.Z deposition is associated with genesis of uterine leiomyoma. Nature (2021). 10.1038/s41586-021-03747-1

9 Greenberg, R. S., Long, H. K., Swigut, T. & Wysocka, J. Single Amino Acid Change Underlies Distinct Roles of H2A.Z Subtypes in Human Syndrome. Cell 178, 1421–1436 e1424 (2019). 10.1016/j.cell.2019.08.002

10 Nacev, B. A. et al. The expanding landscape of ‘oncohistone’ mutations in human cancers. Nature 567, 473–478 (2019). 10.1038/s41586-019-1038-1

11 Truong, D. M. & Boeke, J. D. Resetting the Yeast Epigenome with Human Nucleosomes. Cell 171, 1508–1519 e1513 (2017). 10.1016/j.cell.2017.10.043

12 Talbert, P. B. & Henikoff, S. Histone variants--ancient wrap artists of the epigenome. Nat Rev Mol Cell Biol 11, 264–275 (2010). 10.1038/nrm2861

13 Perino, M. & Veenstra, G. J. Chromatin Control of Developmental Dynamics and Plasticity. Dev Cell 38, 610–620 (2016). 10.1016/j.devcel.2016.08.004

14 Ghosh, R. P. & Meyer, B. J. Spatial Organization of Chromatin: Emergence of Chromatin Structure During Development. Annu Rev Cell Dev Biol 37, 199–232 (2021). 10.1146/annurev-cellbio-032321-035734

15 Rojec, M., Hocher, A., Stevens, K. M., Merkenschlager, M. & Warnecke, T. Chromatinization of Escherichia coli with archaeal histones. Elife 8 (2019). 10.7554/eLife.49038

16 Irwin, N. A. T. & Richards, T. A. Self-assembling viral histones are evolutionary intermediates between archaeal and eukaryotic nucleosomes. Nat Microbiol 9, 1713–1724 (2024). 10.1038/s41564-024-01707-9

17 Kachroo, A. H. et al. Evolution. Systematic humanization of yeast genes reveals conserved functions and genetic modularity. Science 348, 921–925 (2015). 10.1126/science.aaa0769

18 Stargell, L. A. et al. Temporal and spatial association of histone H2A variant hv1 with transcriptionally competent chromatin during nuclear development in Tetrahymena thermophila. Genes Dev 7, 2641–2651 (1993). 10.1101/gad.7.12b.2641

19 Meneghini, M. D., Wu, M. & Madhani, H. D. Conserved histone variant H2A.Z protects euchromatin from the ectopic spread of silent heterochromatin. Cell 112, 725–736 (2003). 10.1016/s0092-8674(03)00123-5

20 Weber, C. M., Ramachandran, S. & Henikoff, S. Nucleosomes are context-specific, H2A.Z-modulated barriers to RNA polymerase. Mol Cell 53, 819–830 (2014). 10.1016/j.molcel.2014.02.014

21 Richter, D. J. et al. EukProt: A database of genome-scale predicted proteins across the diversity of eukaryotes. Peer Community Journal 2 (2022). 10.24072/pcjournal.173

22 Talbert, P. B. et al. A unified phylogeny-based nomenclature for histone variants. Epigenetics Chromatin 5, 7 (2012). 10.1186/1756-8935-5-7

23 Dalmasso, M. C., Sullivan, W. J., Jr. & Angel, S. O. Canonical and variant histones of protozoan parasites. Front Biosci (Landmark Ed) 16, 2086–2105 (2011). 10.2741/3841

24 Heitman, J., Movva, N. R. & Hall, M. N. Targets for cell cycle arrest by the immunosuppressant rapamycin in yeast. Science 253, 905–909 (1991). 10.1126/science.1715094

25 Chernoff, Y. O., Lindquist, S. L., Ono, B., Inge-Vechtomov, S. G. & Liebman, S. W. Role of the chaperone protein Hsp104 in propagation of the yeast prion-like factor [psi+]. Science 268, 880–884 (1995). 10.1126/science.7754373

26 Crowley, P. D. & Gallagher, H. C. Clotrimazole as a pharmaceutical: past, present and future. J Appl Microbiol 117, 611–617 (2014). 10.1111/jam.12554

27 Samakkarn, W., Ratanakhanokchai, K. & Soontorngun, N. Reprogramming of the Ethanol Stress Response in Saccharomyces cerevisiae by the Transcription Factor Znf1 and Its Effect on the Biosynthesis of Glycerol and Ethanol. Appl Environ Microbiol 87, e0058821 (2021). 10.1128/AEM.00588-21

28 Yang, F., Kemp, C. J. & Henikoff, S. Doxorubicin enhances nucleosome turnover around promoters. Curr Biol 23, 782–787 (2013). 10.1016/j.cub.2013.03.043

29 Hoyos-Manchado, R. et al. RNA metabolism is the primary target of formamide in vivo. Scientific reports 7, 15895 (2017). 10.1038/s41598-017-16291-8

30 Torres-Garcia, S. et al. Epigenetic gene silencing by heterochromatin primes fungal resistance. Nature 585, 453–458 (2020). 10.1038/s41586-020-2706-x

31 Luger, K., Mader, A. W., Richmond, R. K., Sargent, D. F. & Richmond, T. J. Crystal structure of the nucleosome core particle at 2.8 A resolution. Nature 389, 251–260 (1997). 10.1038/38444

32 Brewis, H. T. et al. What makes a histone variant a variant: Changing H2A to become H2A.Z. PLoS Genet 17, e1009950 (2021). 10.1371/journal.pgen.1009950

33 Clarkson, M. J., Wells, J. R., Gibson, F., Saint, R. & Tremethick, D. J. Regions of variant histone His2AvD required for Drosophila development. Nature 399, 694–697 (1999). 10.1038/21436

34 Wu, W. H. et al. Swc2 is a widely conserved H2AZ-binding module essential for ATP-dependent histone exchange. Nat Struct Mol Biol 12, 1064–1071 (2005). 10.1038/nsmb1023

35 Luk, E. et al. Stepwise histone replacement by SWR1 requires dual activation with histone H2A.Z and canonical nucleosome. Cell 143, 725–736 (2010). 10.1016/j.cell.2010.10.019

36 Desmoucelles, C., Pinson, B., Saint-Marc, C. & Daignan-Fornier, B. Screening the yeast "disruptome" for mutants affecting resistance to the immunosuppressive drug, mycophenolic acid. J Biol Chem 277, 27036–27044 (2002). 10.1074/jbc.M111433200

37 Judd, J. et al. A rapid, sensitive, scalable method for Precision Run-On sequencing (PRO-seq). bioRxiv (2020). 10.1101/2020.05.18.102277

38 Booth, G. T., Wang, I. X., Cheung, V. G. & Lis, J. T. Divergence of a conserved elongation factor and transcription regulation in budding and fission yeast. Genome Res 26, 799–811 (2016). 10.1101/gr.204578.116

39 Suto, R. K., Clarkson, M. J., Tremethick, D. J. & Luger, K. Crystal structure of a nucleosome core particle containing the variant histone H2A.Z. Nat Struct Biol 7, 1121–1124 (2000). 10.1038/81971

40 Jeronimo, C., Watanabe, S., Kaplan, C. D., Peterson, C. L. & Robert, F. The Histone Chaperones FACT and Spt6 Restrict H2A.Z from Intragenic Locations. Mol Cell 58, 1113–1123 (2015). 10.1016/j.molcel.2015.03.030

41 Narain, A. et al. Targeted protein degradation reveals a direct role of SPT6 in RNAPII elongation and termination. Mol Cell 81, 3110–3127 e3114 (2021). 10.1016/j.molcel.2021.06.016

42 Zumer, K. et al. Two distinct mechanisms of RNA polymerase II elongation stimulation in vivo. Mol Cell 81, 3096–3109 e3098 (2021). 10.1016/j.molcel.2021.05.028

43 Jumper, J. et al. Highly accurate protein structure prediction with AlphaFold. Nature 596, 583–589 (2021). 10.1038/s41586-021-03819-2

44 Harris, M. A. et al. Fission stories: using PomBase to understand Schizosaccharomyces pombe biology. Genetics 220 (2022). 10.1093/genetics/iyab222

45 The Gene Ontology, C. The Gene Ontology Resource: 20 years and still GOing strong. Nucleic Acids Res 47, D330–D338 (2019). 10.1093/nar/gky1055

46 Hong, J. et al. The catalytic subunit of the SWR1 remodeler is a histone chaperone for the H2A.Z-H2B dimer. Mol Cell 53, 498–505 (2014). 10.1016/j.molcel.2014.01.010

47 Kiely, C. M. et al. Spt6 is required for heterochromatic silencing in the fission yeast Schizosaccharomyces pombe. Mol Cell Biol 31, 4193–4204 (2011). 10.1128/MCB.05568-11

48 Fowler, D. M. & Fields, S. Deep mutational scanning: a new style of protein science. Nat Methods 11, 801–807 (2014). 10.1038/nmeth.3027

49 Torres-Garcia, S. et al. SpEDIT: A fast and efficient CRISPR/Cas9 method for fission yeast. Wellcome Open Res 5, 274 (2020). 10.12688/wellcomeopenres.16405.1

50 Vander Heiden, M. G., Cantley, L. C. & Thompson, C. B. Understanding the Warburg effect: the metabolic requirements of cell proliferation. Science 324, 1029–1033 (2009). 10.1126/science.1160809

51 Baldi, S., Bolognesi, A., Meinema, A. C. & Barral, Y. Heat stress promotes longevity in budding yeast by relaxing the confinement of age-promoting factors in the mother cell. Elife 6 (2017). 10.7554/eLife.28329

52 Sanulli, S. et al. HP1 reshapes nucleosome core to promote phase separation of heterochromatin. Nature 575, 390–394 (2019). 10.1038/s41586-019-1669-2

53 Bonisch, C. & Hake, S. B. Histone H2A variants in nucleosomes and chromatin: more or less stable? Nucleic Acids Res 40, 10719–10741 (2012). 10.1093/nar/gks865

54 Chakravarthy, S., Bao, Y., Roberts, V. A., Tremethick, D. & Luger, K. Structural characterization of histone H2A variants. Cold Spring Harb Symp Quant Biol 69, 227–234 (2004). 10.1101/sqb.2004.69.227

55 Osakabe, A. et al. Histone H2A variants confer specific properties to nucleosomes and impact on chromatin accessibility. Nucleic Acids Res 46, 7675–7685 (2018). 10.1093/nar/gky540

56 Arimura, Y. et al. Structural basis of a nucleosome containing histone H2A.B/H2A.Bbd that transiently associates with reorganized chromatin. Scientific reports 3, 3510 (2013). 10.1038/srep03510

57 Horikoshi, N., Arimura, Y., Taguchi, H. & Kurumizaka, H. Crystal structures of heterotypic nucleosomes containing histones H2A.Z and H2A. Open Biol 6 (2016). 10.1098/rsob.160127

58 Kohestani, H. & Wereszczynski, J. Effects of H2A.B incorporation on nucleosome structures and dynamics. Biophys J 120, 1498–1509 (2021). 10.1016/j.bpj.2021.01.036

59 Horikoshi, N., Kujirai, T., Sato, K., Kimura, H. & Kurumizaka, H. Structure-based design of an H2A.Z.1 mutant stabilizing a nucleosome in vitro and in vivo. Biochem Biophys Res Commun 515, 719–724 (2019). 10.1016/j.bbrc.2019.06.012

60 Bagert, J. D. et al. Oncohistone mutations enhance chromatin remodeling and alter cell fates. Nat Chem Biol 17, 403–411 (2021). 10.1038/s41589-021-00738-1

61 Schwartzentruber, J. et al. Driver mutations in histone H3.3 and chromatin remodelling genes in paediatric glioblastoma. Nature 482, 226–231 (2012). 10.1038/nature10833

62 Lu, C. et al. Histone H3K36 mutations promote sarcomagenesis through altered histone methylation landscape. Science 352, 844–849 (2016). 10.1126/science.aac7272

63 Eddy, S. R. Accelerated Profile HMM Searches. PLoS Comput Biol 7, e1002195 (2011). 10.1371/journal.pcbi.1002195

64 Draizen, E. J. et al. HistoneDB 2.0: a histone database with variants--an integrated resource to explore histones and their variants. Database (Oxford) 2016 (2016). 10.1093/database/baw014

65 Katoh, K. & Standley, D. M. MAFFT multiple sequence alignment software version 7: improvements in performance and usability. Mol Biol Evol 30, 772–780 (2013). 10.1093/molbev/mst010

66 Darriba, D. et al. ModelTest-NG: A New and Scalable Tool for the Selection of DNA and Protein Evolutionary Models. Mol Biol Evol 37, 291–294 (2020). 10.1093/molbev/msz189

67 Kozlov, A. M., Darriba, D., Flouri, T., Morel, B. & Stamatakis, A. RAxML-NG: a fast, scalable and user-friendly tool for maximum likelihood phylogenetic inference. Bioinformatics 35, 4453–4455 (2019). 10.1093/bioinformatics/btz305

68 Letunic, I. & Bork, P. Interactive Tree Of Life (iTOL) v4: recent updates and new developments. Nucleic Acids Res 47, W256–W259 (2019). 10.1093/nar/gkz239

69 Maundrell, K. nmt1 of fission yeast. A highly transcribed gene completely repressed by thiamine. Journal of Biological Chemistry 265, 10857–10864 (1990). 10.1016/s0021-9258(19)38525-4

70 Yeast Extract with Supplements (YES). Cold Spring Harbor Protocols 2016 (2016). 10.1101/pdb.rec091355

71 Bahler, J. et al. Heterologous modules for efficient and versatile PCR-based gene targeting in Schizosaccharomyces pombe. Yeast 14, 943–951 (1998). 10.1002/(SICI)1097-0061(199807)14:10<943::AID-YEA292>3.0.CO;2-Y

72 Murray, J. M., Watson, A. T. & Carr, A. M. Transformation of Schizosaccharomyces pombe: Lithium Acetate/ Dimethyl Sulfoxide Procedure. Cold Spring Harb Protoc 2016, pdb prot090969 (2016). 10.1101/pdb.prot090969

73 Gibson, D. G. et al. Enzymatic assembly of DNA molecules up to several hundred kilobases. Nat Methods 6, 343–345 (2009). 10.1038/nmeth.1318

74 Grimm, C., Kohli, J., Murray, J. & Maundrell, K. Genetic engineering of Schizosaccharomyces pombe: a system for gene disruption and replacement using the ura4 gene as a selectable marker. Mol Gen Genet 215, 81–86 (1988). 10.1007/BF00331307

75 Lowary, P. T. & Widom, J. New DNA sequence rules for high affinity binding to histone octamer and sequence-directed nucleosome positioning. J Mol Biol 276, 19–42 (1998). 10.1006/jmbi.1997.1494

76 Tachiwana, H. et al. Structural basis of instability of the nucleosome containing a testis-specific histone variant, human H3T. Proc Natl Acad Sci U S A 107, 10454–10459 (2010). 10.1073/pnas.1003064107

77 Taguchi, H., Horikoshi, N., Arimura, Y. & Kurumizaka, H. A method for evaluating nucleosome stability with a protein-binding fluorescent dye. Methods 70, 119–126 (2014). 10.1016/j.ymeth.2014.08.019

78 DeAngelis, M. M., Wang, D. G. & Hawkins, T. L. Solid-phase reversible immobilization for the isolation of PCR products. Nucleic Acids Res 23, 4742–4743 (1995). 10.1093/nar/23.22.4742

79 Bray, N. L., Pimentel, H., Melsted, P. & Pachter, L. Near-optimal probabilistic RNA-seq quantification. Nat Biotechnol 34, 525–527 (2016). 10.1038/nbt.3519

80 Love, M. I., Huber, W. & Anders, S. Moderated estimation of fold change and dispersion for RNA-seq data with DESeq2. Genome Biol 15, 550 (2014). 10.1186/s13059-014-0550-8

81 Ramírez, F. et al. deepTools2: a next generation web server for deep-sequencing data analysis. Nucleic Acids Res. 44, W160–W165 (2016). 10.1093/nar/gkw257

82 Mirdita, M. et al. ColabFold: making protein folding accessible to all. Nat Methods 19, 679–682 (2022). 10.1038/s41592-022-01488-1

